# Hippo signaling restricts cells in the second heart field that differentiate into Islet-1-positive atrial cardiomyocytes

**DOI:** 10.1101/154765

**Authors:** Hajime Fukui, Takahiro Miyazaki, Hiroyuki Ishikawa, Hiroyuki Nakajima, Naoki Mochizuki

**Affiliations:** Department of Cell Biology, National Cerebral and Cardiovascular Center Research Institute, Fujishirodai 5-7-1, Suita, Osaka 565-8565, Japan; AMED-CREST

## Abstract

Cardiac precursor cells (CPCs) in the first heart field (FHF) and the second heart field (SHF) present at both arterial and venous poles assemble to form a cardiac tube in zebrafish. Hippo kinase cascade is essential for proper heart formation; however, it remains elusive how Hippo signal contributes to early cardiac fate determination. We here demonstrate that mutants of *large tumor suppressor kinase 1/2* (*lats1/2*) exhibited an increase in a SHF marker, Islet1 (Isl1)-positive and *hand2* promoter-activated venous pole atrial cardiomyocytes (CMs) and that those showed expansion of the domain between between the anterior and the posterior lateral plate mesoderm. Consistently, TEAD-8 dependent transcription was activated in caudal region of the left ALPM cells that gave rise to the venous pole atrial CMs. Yap1/Wwtr1-promoted *bmp2b* expression was essential for Smad-regulated *hand2* expression in the left ALPM, indicating that Hippo signaling restricts the SHF cells originating from the left ALPM that move toward the venous pole.

## Introduction

Heart forms mainly according to the assembly of cardiomyocytes (CMs) and blood vessel-constituting cells that originate from anterior lateral plate mesoderm (ALPM). The ALPM can be distinguished from the posterior lateral plate mesoderm (PLPM) at the 6-8 somite stage (ss) (Waxman et al., 2008). The embryonic heart field is specified during formation of ALPM (Fishman and Chien, 1997). Bilaterally located cells that finally differentiate into heart field cells migrate medially and fuse at the midline to form the cardiac tube (Staudt and Stainier, 2012). Several signaling pathways exert roles to restrict the heart field potency at the both rostral and caudal boundaries of the ALPM. The retinoic acid (RA) signaling determines the forelimb field by restricting the posterior end of heart field (Waxman et al., 2008). Tal1 and Etv2 are transcriptional factors for vascular and hematopoietic lineage specification, respectively, and repress cardiac specification in rostral ALPM region (Schoenebeck et al., 2007). While the heart field is defined by restriction of the other organ fields in the ALPM (Fishman and Chien, 1997), it remains unclear which signals regulate the differentiation of ALPM into heart field and the other fields.

In vertebrates, the first heart field (FHF)-derived CMs and the second heart field (SHF)-derived CMs contribute to the initial tube formation and to the accretion of CMs at the arterial and venous poles, respectively (Kelly et al., 2014). These cells in the FHF and SHF are thus cardiac precursor cells (CPCs). During heart development in chick and mouse embryos, the progenitors of the venous pole are located most laterally and caudally in the ALPM (Abu-Issa and Kirby, 2008; Galli et al., 2008). While mammalian SHF-derived CMs have been characterized extensively using lineage-tracing (Cai et al., 2003; Galli et al., 2008; Abu-Issa and Kirby, 2008), specification and expansion of zebrafish SHF-1 derived cells remains unclear. Successive phases of CM differentiation are conserved among vertebrates during the development of myocardium (Staudt and Stainier, 2012), although zebrafish heart consists of one atrium and one ventricle in contrast to four-chambered heart in mammals.

In zebrafish, a LIM homeodomain transcription factor, Islet-1 (Isl1)-positive SHF cells give rise to CMs in the venous pole and consequent inflow tract (IFT) CMs of the atrium, whereas Isl2b and latent TGFβ binding protein 3 (Ltbp3)-positive SHF cells become CMs only at the arterial pole and subsequently contribute to the formation of outflow tract (OFT) of the ventricle (de Pater et al., 2009; Zhou et al., 2011; Witzel et al., 2017). While the number of CMs in the venous pole in *isl1* mutants is decreased, that at the arterial poles remains unchanged (de Pater et al., 2009). Ltbp3-positive SHF cells express transcription factors, Nkx2.5 and Mef2c that are also expressed in the FHF (Guner-Ataman et al., 2013; Hinits et al., 2012). Another transcription factor, Hand2, a basic helix-loop-helix transcription factor, is involved in the arterial pole formation by the CPC in the SHF (Schindler et al., 2014). While the essential and potential roles of these transcription factors during cardiogenesis have been reported (Guner-Ataman et al., 2013; Hinits et al., 2012; Schindler et al., 2014), it remains elusive how the expression of these transcription factors is regulated.

Hippo signaling pathway defines the number of cells in tissue/organ including heart (Zhou et al., 2015). Mammalian Ste20-like serine/threonine kinase 1 and 2 (Mst1/2, mammalian orthologs of fruit fly, Hippo) phosphorylate Large tumor suppressor kinase 1 and 2 (Lats1/2). Phosphorylated Lats1/2 induce nuclear export of Yes-associated protein 1 (Yap1) / WW domain containing transcription regulator 1 (Wwtr1, also known as Taz), thereby inhibiting Yap1/Wwtr1-TEA domain (TEAD) transcription factor complex-8 dependent expression of genes essential for cell specification, proliferation, survival, and differentiation (Zhao et al., 2008; Nishioka et al., 2009). Hippo signaling has been implicated in heart formation as well as regeneration after myocardial injury (Lin et al., 2014; von Gise et al., 2012; Xin et al., 2013). Nuclear Yap1 drives the proliferation of CM in adult and fetal mouse heart. Mice depleted of *Lats2*, *Salvador* (*Salv*), or *Mst1/2* using CM-specific Cre drivers exhibit a hypertrophic growth owing to an increase of CMs (Zhou et al., 2015). *Yap1* and *Wwtr1* double-null mutant mice are embryonic lethal before the blastula stage (Nishioka et al., 2009), suggesting the essential role for Yap1/Wwtr1 in early cardiovascular development, although it is unclear whether Yap1/Wwtr1 function in the FHF/SHF-dependent cardiac development.

In this study, we demonstrate that Lats1/2-Yap1/Wwtr1-regulated hippo signaling determines the fate of cells between ALPM and PLPM that influences the number of both *hand2* and *isl1* promoter-activated SHF cells. Moreover, we reveal that Yap1/Wwtr1 promote *bone morphogenetic protein-2b* (*bmp2b*) expression in the ALPM required for accretion in the venous pole and subsequent inflow tract (IFT) atrial CMs development.

## Results

### Lats1/2 are involved in atrial CMs development

To examine whether Yap1/Wwtr1-dependent transcription determines the CMs number during early cardiogenesis, we developed *lats1* and *lats2* knockout (KO) fish using transcription activator-like effector nuclease (TALEN) techniques. The fish with *lats1^ncv107^* and *lats2^ncv108^* allele lacked 10 bp at Exon 2 and 16 bp at Exon 3, respectively, and had premature stop codons due to frame shifts (Figure 1—Figure supplement 1A). Either *lats1^ncv107^* KO fish or *lats2^ncv108^* KO fish was viable with no apparent defect (data not shown). However, almost all of the *lats1^ncv107^lats2^ncv108^* double KO (*lats1/2* DKO) larvae died before 15 days post-fertilization (dpf) (Figure 1—Figure supplement 1B). We examined the effect of Lats1/2 depletion on heart development by counting CM numbers in atrium and ventricle in *Tg(myosin heavy chain 6 [myh6]:Nls-tdEosFP);Tg(myl7:Nls-mCherry)* larvae with *lats1/2* mutant alleles that expressed Nls-tagged tandem EOS fluorescent protein under the control of atrium-specific *myh6* promoter and Nls-mCherry under the control of *myl7* promoter (Figure 1A). The number of atrial CMs but not ventricular CMs was significantly increased in *lats1^wt/ ncv107^lats2^ncv108^* embryos and *lats1^ncv107^lats2^ncv108^* embryos at 74 hours post-fertilization (hpf) (Figure 1B, C and Figure 1—source data 1).

**Figure 1.**
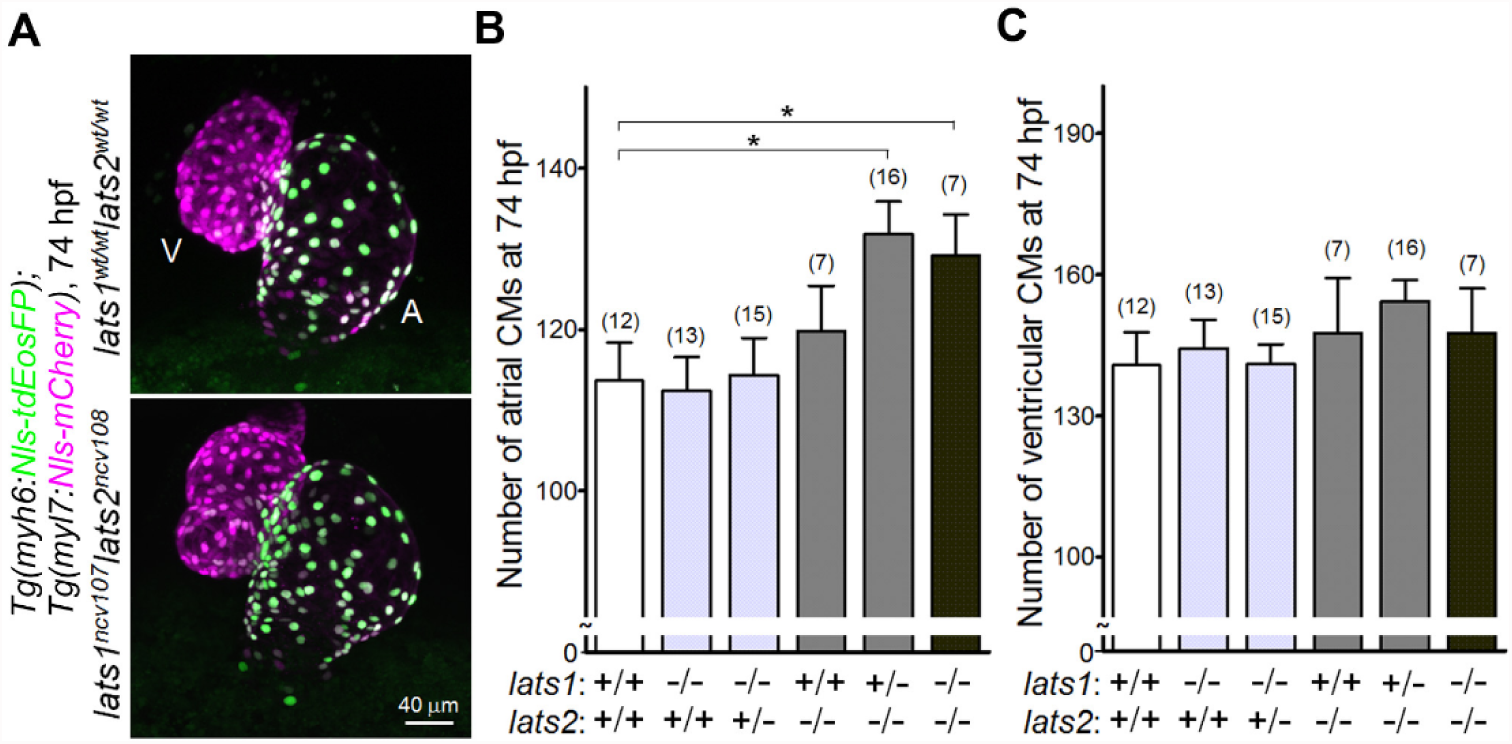
Knockout of *lats1/2* genes leads to an increase of the number of atrial, but not ventricular CMs during early development. **(A)** 3D confocal stack images of *Tg*(*myh6:Nls-tdEosFP);Tg(myl7:Nls-mCherry*) embryos at 74 hpf of the *lats1^wt/wt^lats2^wt/wt^* (top) and *the lats1^ncv107^lats2^ncv108^* (bottom). Atrial (A) and ventricular (V) cardiomyocytes (CMs) are EosFP-4 positive cells and EosFP-negative mCherry-positive cells, respectively. Ventral view, anterior to the top. The confocal 3D-stack images are a set of representative images of eight independent experiments. **(B, C)** Quantitative analyses of the number of atrial **(B)** and ventricular **(C)** CMs of the embryos at 74 hpf with alleles indicated at the bottom. Plus (+) and minus (-) indicate the *wt* allele and the allele of *ncv107* or *ncv108* in *lats1* or *lats2* genes, respectively. In the following graphs, total number of larvae examined in the experiment is indicated on the top of column unless otherwise described. ^*^p < 0.05.

To examine whether Yap1/Wwtr1-dependent transcription is activated during embryogenesis and in CMs, we used two Tead reporter Tg lines: one expressed human TEAD2 lacking amino-terminus (1–113 aa) fused with GAL4 DNA-binding domain under the control of *eukaryotic translation elongation factor 1 alpha 1, like 1* (*eef1a1l1*) promoter, *Tg(eef1a1l1:galdb-hTEAD2*Δ*N-2A-mCherry)* ; the other expressed that under the control of CM-specific *myosin light polypeptide 7* (*myl7*) promoter, *Tg(myl7:galdb-hTEAD2*Δ*N-2A-mCherry).* In these Tg fish crossed with *Tg(uas:GFP)*, when Yap1 or Wwtr1 entered the nuclei, GFP expression was promoted according to the Gal4-UAS system (Fukui et al., 2014). Hereafter, we named the former, general Tead reporter, and the latter, CM Tead reporter, respectively. GFP-expressing cells reflected the nuclear translocation of Yap1 and/or Wwtr1. Yap1/Wwtr1-dependent transcriptional activation defined by increased GFP expression was found in *lats1/2* DKO embryos and *lats1/2* morphants (Figure 1—Figure supplement 2A), suggesting that these KO fish allow us to examine when and where Yap1/Wwtr1-Tead complex-dependent transcription was activated. The CPCs in the venous pole become the IFT atrial CMs (de Pater et al., 2009). We confirmed Yap1/Wwtr1-dependent transcription in IFT atrial CMs by the CM Tead reporter fish embryos at 74 hpf (Figure 1—Figure supplement 2B). At 24 hpf, GFP expression was observed in the *myl7* promoter-activated CMs of the venous pole but not the other area of general Tead reporter crossed with *Tg(myl7:Nuclear localization signal [Nls]-tagged mCherry)* fish (Figure 1— Figure supplement 2C). In this Tg fish embryos, mCherry expression driven by *eef1a1l1* and 2A peptide was subtle compared to that driven by *myl7* promoter. These data suggest that Lats1/2 restrict the Yap1/wwtr1-Tead signal-dependent increase in atrial CM number during early cardiac development.

### Lats1/2 determine the number of CMs derived from the *hand2* promoter-activated CMs

We assumed that an increase in atrial CMs might be ascribed to an increase in CPCs in the SHF, because Yap1/Wwtr1-dependent transcription was observed in the venous pole of heart tube and the IFT CMs of atrium (Figure 1—Figure supplement 2B, C). Firstly, we examined the effect of nuclear Yap1/Wwtr1 on early cardiogenesis by investigating the expression of transcription factors, *nkx2.5*, *hand2*, and *gata4*, essential for early CPC differentiation (Schoenebeck et al., 2007). Among these transcription factors, *hand2* mRNA expression was significantly up-regulated in the *lats1/2* morphants (Figure 2A and Figure 2— source data 1). The expression of *nkx2.5* and *gata4* mRNAs was unaffected by the depletion of Lats1/2 (Figure 2A and Figure 2—source data 1). The whole mount *in situ* hybridization (WISH) analyses revealed that *hand2* expression was increased in the domain that was supposed to give rise to the heart in *lats1^wt/ ncv107^lats2^ncv108^* embryos, *lats1/2* DKO embryos and *lats1/2* morphants at 22 hpf (Figure 2B and Figure 2—Figure supplement 1A). These data suggest that Lats1/2 might determine the number of atrial CMs through the control of *hand2* expression.

**Figure 2.**
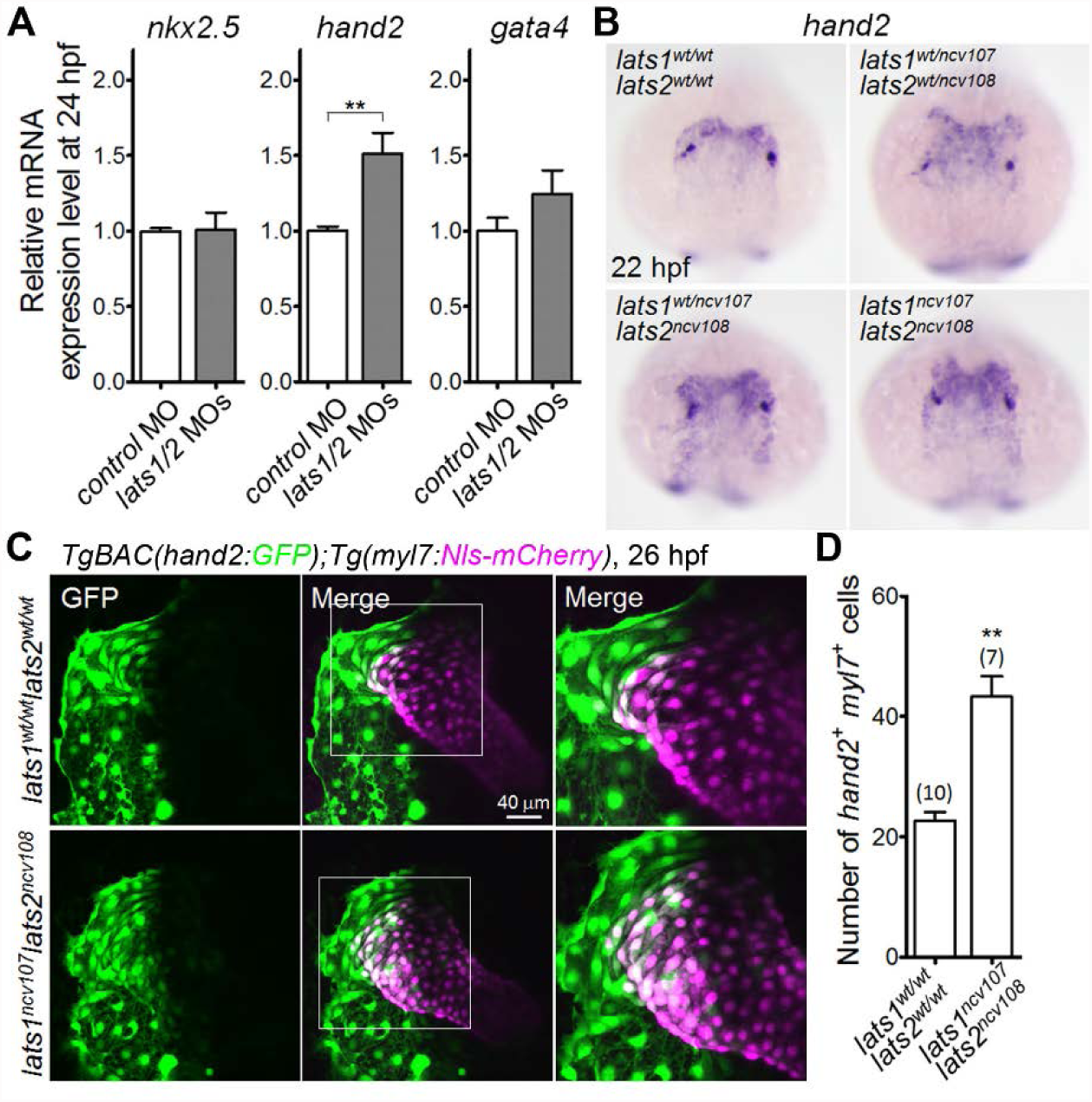
Knockout of *lats1/2* results in an increase in the both *myl7* and *hand2* promoter-activated cells in the venous pole. **(A)** Quantitative-PCR analyses of expression of *nkx2.5*, *hand2*, and *gata4* mRNAs in the whole embryos at 24 hpf injected with the morpholino (MO) indicated at the bottom (n=4). Relative expression of mRNA in the morphants to that of the control is calculated. **(B)** Whole mount in situ hybridization (WISH) analyses of the embryos at 22 hpf of the *lats1^wt/wt^lats2^wt/wt^*, *lats1^wt/ncv107^lats2^wt/ncv108^*, *lats1^wt/ncv107^lats2^ncv108^* and the *lats1^ncv107^lats2^ncv108^* indicated at the top using antisense probe for *hand2*. **(C)** 3D confocal stack images of the *TgBAC*(*hand2:GFP);Tg(myl7:Nls-mCherry*) embryos of the *lats1^wt/wt^lats2^wt/wt^* (upper panels) and *the lats1^ncv107^lats2^ncv108^* (bottom panels) at 26 hpf. GFP images (left), merged images of GFP image and mCherry image (center), and enlarged images of boxed regions in the center panels (right). **(D)** Quantitative analysis of the number of both *hand2* and *myl7* promoter-activated cells at 26 hpf. All images in Figure 2 are dorsal view, anterior to the top. The confocal 3D-stack images and ISH images are a set of representative images of at least four independent experiments.^**^p < 0.01.

Overexpression of Hand2 increases the number of SHF-derived CMs but not FHF-derived CMs in zebrafish (Schindler et al., 2014). To investigate the relevance of *hand2* expression to CM number in the atrium, we tried to count the number of *hand2*-positive CMs at early time point using Tg fish expressing GFP under the control of *hand2* BAC promoter; *TgBAC(hand2:GFP) (Yin et al., 2010)*. We crossed this Tg fish with *Tg(myl7:Nls-mCherry)* to count the number of both *hand2* and *myl7* promoter-activated CMs at 26 hpf. Both promoter-activated CMs were localized at the anterior side of growing heart tube corresponding to the venous pole (Figure 2C). In the *lats1/2* DKO embryos, the number of both promoter-activated CMs was significantly increased in the venous pole (Figure 2C, D and Figure 2—source data 1). Consistent with these results, the both promoter-activated cells were increased in the venous pole in the *lats1/2* morphants (Figure 2—Figure supplement 1B). To examine the function of Yap1/Wwtr1, we developed double KO of *yap1* (Figure 2—Figure supplement 2A) and *wwtr1* (Nakajima et al., 2017). In the *yap1* and *wwtr1* DKO embryos, *hand2* promoter-activated cells were greatly reduced. Those double mutant embryos exhibited cardia bifida (Figure 2—Figure supplement 2B). These results suggest that Lats1/2 are involved in the formation of *hand2* promoter-activated cells present in the venous pole cells that give rise to the IFT atrial CMs.

### *hand2* promoter-activated cells in the caudal end of the left ALPM migrate toward venous pole of heart tube

The progenitors of the venous pole are located most caudally in the ALPM in the mammalian heart (Abu-Issa and Kirby, 2008; Galli et al., 2008). To investigate how *hand2* promoter-activated cells contribute the venous pole cells that differentiate into CPCs, we time-lapse imaged them from 14 hpf to 26 hpf. Certain number of cells in the caudal side of the left ALPM migrated toward the venous pole, whereas those of the right ALPM migrate into the arterial pole (Figure 3A, B, and Video 1). The cell tracking analyses demonstrated that both former and latter cells moved toward the region where the cardiac disk developed at 20 hpf and subsequently became venous pole and arterial pole, respectively (Figure 3A-C). These results indicate that *hand2* promoter-activated cells in the caudal region of the left ALPM differentiate into the venous pole cells. This directional migration of the *hand2* promoter-activated cells in the caudal region of the ALPM cells was further confirmed by the embryos with situs inversus. The *polycystin-2* (*pkd2)* morphant causes the randomization of left-8 right patterning which often results in situs inversus (Bisgrove et al., 2005). The caudal region of right ALPM moved toward the venous pole cells of the heart of the embryos with situs inversus. (Figure 3—Figure supplement 1 and Video 2). Collectively, the directional migration of the cells in the caudal region of the ALPM cells toward the venous pole might be predetermined by uncertain signaling.

**Figure 3.**
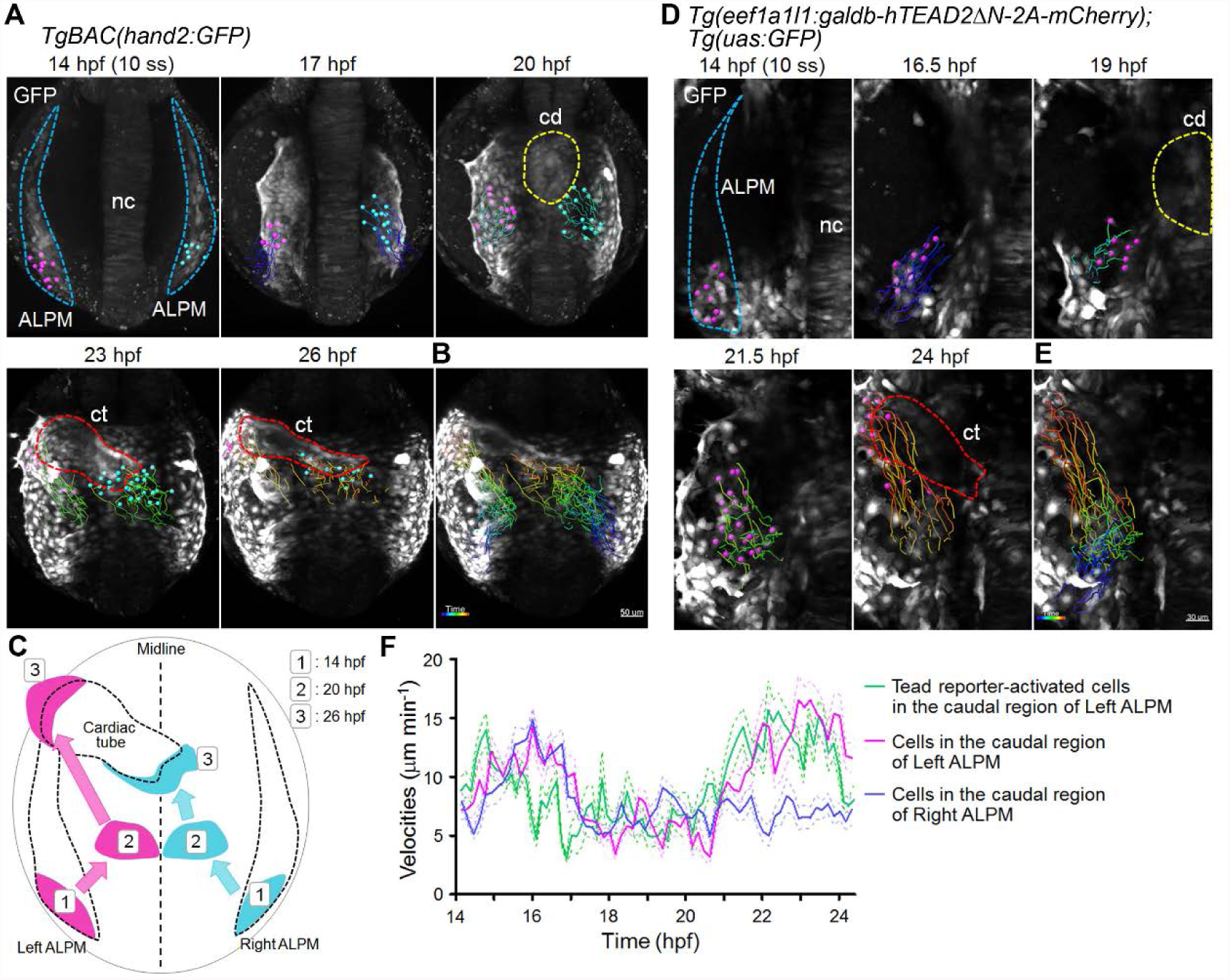
Tead transcription-activated cells in the caudal region of the left ALPM move to venous pole **(A, B)** Time-sequential 3D-rendered confocal images of a *TgBAC*(*hand2:GFP*) embryo from 14 hpf (10 ss) to 26 hpf as indicate at the top. Spots of magenta and cyan denote the cells in the caudal part of left and right ALPM, respectively. **(B)** Tracking of caudal end *hand2* promoter-activated ALPM cells from 14 hpf to 26 hpf. The color of the tracks changes from blue to red according to the time after imaging (0 h to 12 h). Notochord, nc; cardiac disc, cd; cardiac tube, ct. ALPM, cd, and ct are marked by the blue, yellow, and red broken lines, respectively. **(C)** Schematic illustration of trajectory patterns of the caudal end ALPM cells from 14 hpf to 26 hpf. Magenta and cyan denote the region of caudal region of left and right ALPM, respectively. The cells in the caudal region of left (magenta) and right (cyan) ALPM moved from the region 1 (14 hpf) to the region 3 (26 hpf) through the region 2 (20 hpf). **(D, E)** Time-sequential 3D-rendered confocal images of a *Tg(eef1a1l1:galdb-hTEAD2*Δ*N-2A-mCherry);Tg(uas:GFP*) embryo from 14 hpf (10 ss) to 24 hpf. Spots of magenta denote the Tead reporter-activated cells in the caudal region of the left ALPM. **(E)** Tracking of Tead reporter-activated cells from 14 hpf to 24 hpf. The color of the tracks changes from blue to red according to the time after imaging (0 h to 10 h). **(F)** Mean velocities of movement of *hand2* promoter-activated cells in the caudal region of left (magenta) and right (blue) ALPM and that of Tead reporter-activated cells in the caudal region of the left ALPM (green) from 14 hpf to 24 hpf. Broken lines indicate the SEM of the mean line. All images in Figure 3 are dorsal view, anterior to the top. The confocal 3D-stack images are a set of representative images of six independent experiments.

We next tracked Tead transcription-activated cells using general Tead reporter fish embryos. At 14 hpf, the Tead reporter-activated cells were located in the caudal region of the left ALPM (Figure 3D). Subsequently, these cells moved toward the venous pole similarly to the *hand2* promoter-activated cells (Figure 3D, E, and Video 3). We further found that the migration speed of the cells initially located in the caudal region of the left ALPM (the pink line of Figure 3F and Figure 3—source data 1) migrating toward the venous pole was very similar to that of the Tead reporter-activated cells (from 21 hpf to 24 hpf) (the green line of Figure 3F and Figure 3—source data 1), suggesting that those cells are likely to be the same cells and that Tead-activated cells become the venous pole cells. Furthermore, these data imply that the number of the cells might be decided by Hippo signaling.

### Lats1/2 determine the number of IFT CMs derived from the Isl1-positive SHF cells

We, next, investigated whether both *hand2* and *myl7* promoter-activated cells are the SHF-derived cells that add to the growing heart tube in the venous pole. We used Isl1 as a SHF marker, because Isl1 plays an essential role for the development of CPCs in the SHF and forms complex with key regulatory molecules for SHF development, such as Hand2 (Cai et al., 2003; Caputo et al., 2015). We found that in the *lats1/2* DKO embryos, Isl1-positive SHF cells were increased and overlapped both *hand2* and *myl7* promoter-activated CMs in the very left-rostral end of heart tube (Figure 4A, brackets). Consistently, the number of both promoter-activated Isl1-positive CMs in the *lats1/2* morphants was increased (Figure 4—Figure supplement 1A). Furthermore, general Tead reporter-activated cells were positive for Isl1 in the venous pole and were increased by the depletion of Lats1/2 (Figure 4—Figure supplement 1B, arrows and Figure 4—Figure supplement 1C).

**Figure 4.**
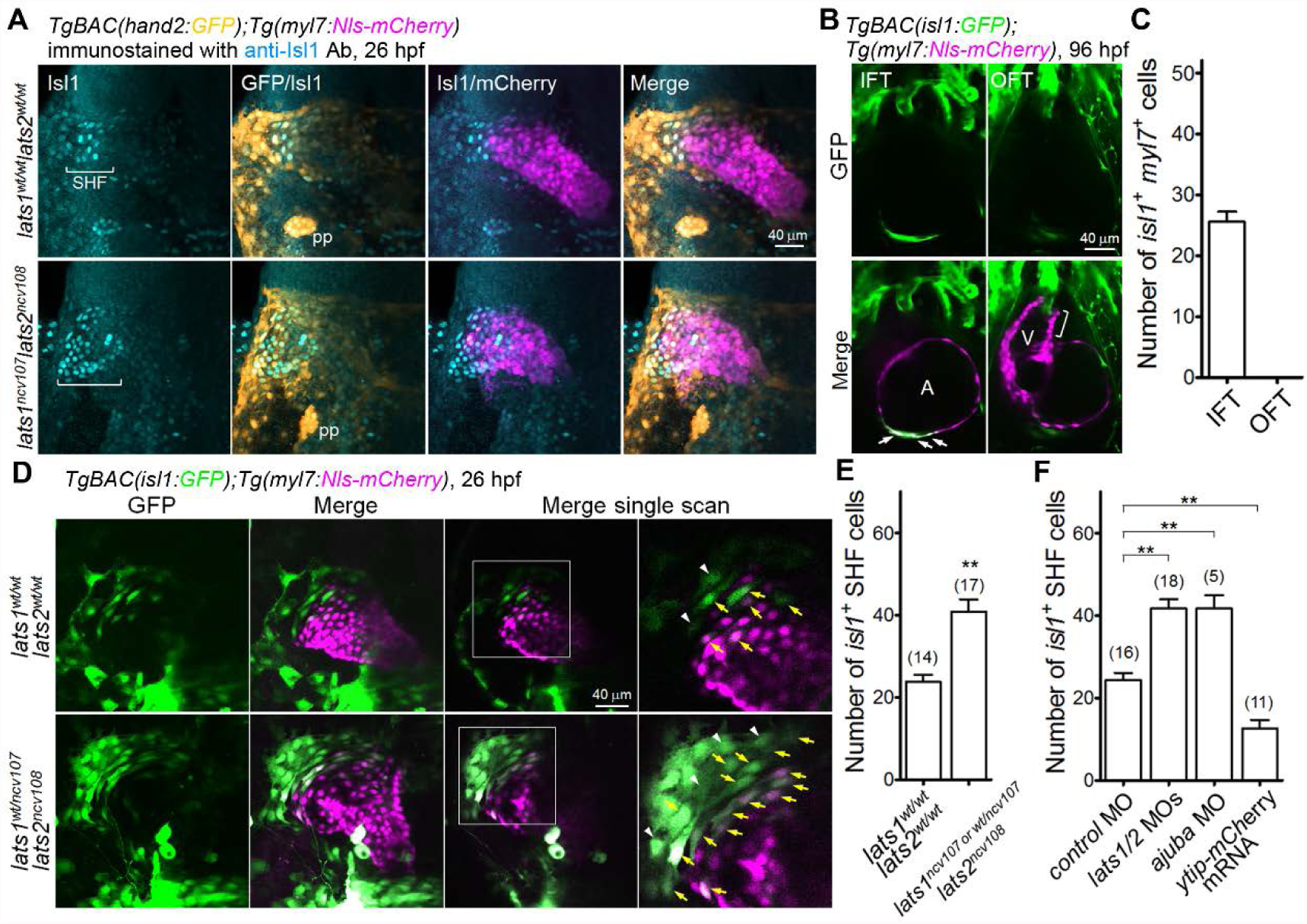
Hippo signaling is involved in the formation of the Isl1-positive SHF cells in the venous pole. **(A)** 3D confocal stack images of the *TgBAC*(*hand2:GFP);Tg(myl7:Nls-mCherry*) embryos with *lats1^wt/wt^lats2^wt/wt^* allele (upper panels) and *lats1^ncv107^lats2^ncv108^* allele (bottom panels) immunostained with anti-Isl1 antibody (anti-Isl1 Ab) at 26 hpf. Brackets denote the SHF cells that are Isl1-positive and both *hand2* and *myl7* promoter-activated cells and that are Isl1-positive and *hand2* promoter-activated cells in contact with *myl7* promoter-activated cells. pp indicates the pharyngeal pouch that expresses *hand2* promoter-activated GFP signal. Dorsal view, anterior to the top. The first, second, third and fourth images are Isl-1 immunostaining, the merged image of GFP image and Isl1 immunostaining, the merged image of Isl1 immunostaining and mCherry image, and the merged of all the three (GFP, mCherry, and Isl1 immunostaining), respectively. **(B)** Single-scan confocal images of the *TgBAC*(*isl1:GFP);Tg(myl7:Nls-mCherry*) embryos at 96 hpf. Both *isl1* and *myl7* promoter-activated cells are in the inflow tract (IFT) cells (arrows) but not in the outflow tract (OFT) cells (bracket). A, atrium; V, ventricle. Ventral view, anterior to the top. The confocal images are a set of representative images of four independent experiments. **(C)** Quantitative analyses of the number of both *isl1* and *myl7* promoter-activated cells in the IFT and OFT at 96 hpf (n=10). **(D)** Confocal images of the *TgBAC*(*isl1:GFP);Tg(myl7:Nls-mCherry*) embryos of the *lats1^wt/wt^lats2^wt/wt^* and the *lats1^wt/ncv107^lats2^ncv108^* at 26 hpf. The boxed regions are enlarged in the most right panels. Yellow arrows indicate both *isl1* and *myl7* promoter-activated cells in the venous pole. White arrowheads indicate the *isl1* promoter-activated cells in contact with *myl7* promoter-activated cells. 3D confocal stack images (left two panels) and single-scan images (right two panels). Dorsal view, anterior to the top. **(E, F)** Quantitative analyses of the number of the *isl1* promoter-activated SHF cells in the venous pole of *lats1^wt/wt^lats2^wt/wt^* embryos and either *lats1^wt/ncv107^lats2^ncv108^* embryos or *lats1/2* DKO embryos **(E)**, and embryos shown in Figure 4— Figure supplement 1E **(F)**. Both *isl1* and *myl7* promoter-activated cells and *isl1* promoter-1 activated cells in contact with *myl7* promoter-activated cell were counted as SHF cells. The confocal 3D-stack images and single-scan (2 μm) images are a set of representative images of at least four independent experiments. ^**^p < 0.01.

To characterize *isl1* promoter-activated SHF cells during early stage and to examine whether those cells become CMs of the IFT, we generated Tg fish expressing GFP under the control of *isl1* BAC promoter; *TgBAC(isl1:GFP)*. *isl1* promoter-activated cells were found in the IFT of atrium at 4 dpf (Figure 4B, C and Figure 4—source data 1). While *isl1* promoter-activated cells were observed in the endocardium and epicardium at 96 hpf (Figure 4—Figure supplement 1D, arrows and arrowheads), those cells were found in neither arterial pole, OFT, nor ventricular myocardium until 4 dpf (Figure 4B, C and Figure 4—source data 1). The number of *isl1* promoter-activated cells in the venous pole was significantly increased in either *lats1^wt/ncv107^lats2^ncv108^* embryos or *lats1/2* DKO embryos at 26 hpf (Figure 4D, E and Figure 4—source data 1). Consistent with this, the *isl1* promoter-activated SHF cells were significantly increased in the venous pole in the *lats1/2* morphants (Figure 4F, Figure 4— Figure supplement 1E and Figure 4—source data 1). Ajuba, a LIM-domain family protein, restricts Isl1-positive SHF cells by binding to Isl1 (Witzel et al., 2012). The number of *isl1* promoter-activated SHF cells were increased in the *ajuba* morphants (Figure 4F, Figure 4—Figure supplement 1E and Figure 4— source data 1). We also found that in the embryos expressing a mCherry-0 tagged dominant-negative form of Yap1/Wwtr1-Tead-dependent transcription (ytip-mCherry) (Fukui et al., 2014), the *isl1* promoter-activated SHF cells were significantly decreased in the venous pole (Figure 4F, Figure 4—Figure supplement 1E and Figure 4—source data 1). Therefore, we confirmed that Lats1/2-mediated hippo signaling is involved in the accretion of SHF-derived CPCs in the venous pole.

### Yap1/Wwtr1 promote the differentiation of SHF cells from the caudal end of ALPM

Tead reporter activation in the cells of ALPM as shown in Figure 3F prompted us to ask whether Lats1/2-Yap1/Wwtr1 signal is involved in either or both proliferation and/or specification of those cells into the Isl1-positive SHF cells from the ALPM. To investigate whether the increase in the number of SHF cells in *lats1/2* DKO embryos and *lats1/2* morphants is ascribed to the proliferation of the SHF cells that have differentiated from the ALPM, we examined proliferation of *isl1* promoter-activated cells by the EdU incorporation assay. The number of *isl1* promoter-activated EdU-positive CM of the *lats1/2* morphants was comparable to that of the control (Figure 5A, B). There was no difference of the number of EdU-positive blood cells and endocardial cells among the two groups (data not shown). Furthermore, the timing of EdU incorporation did not affect the results of the proliferation analyses (Figure 5B), suggesting that the increased number of *isl1*-positive SHF cells in the depletion of Lats1/2 is not caused by the cell proliferation after the differentiation of SHF cells from the ALPM.

**Figure 5.**
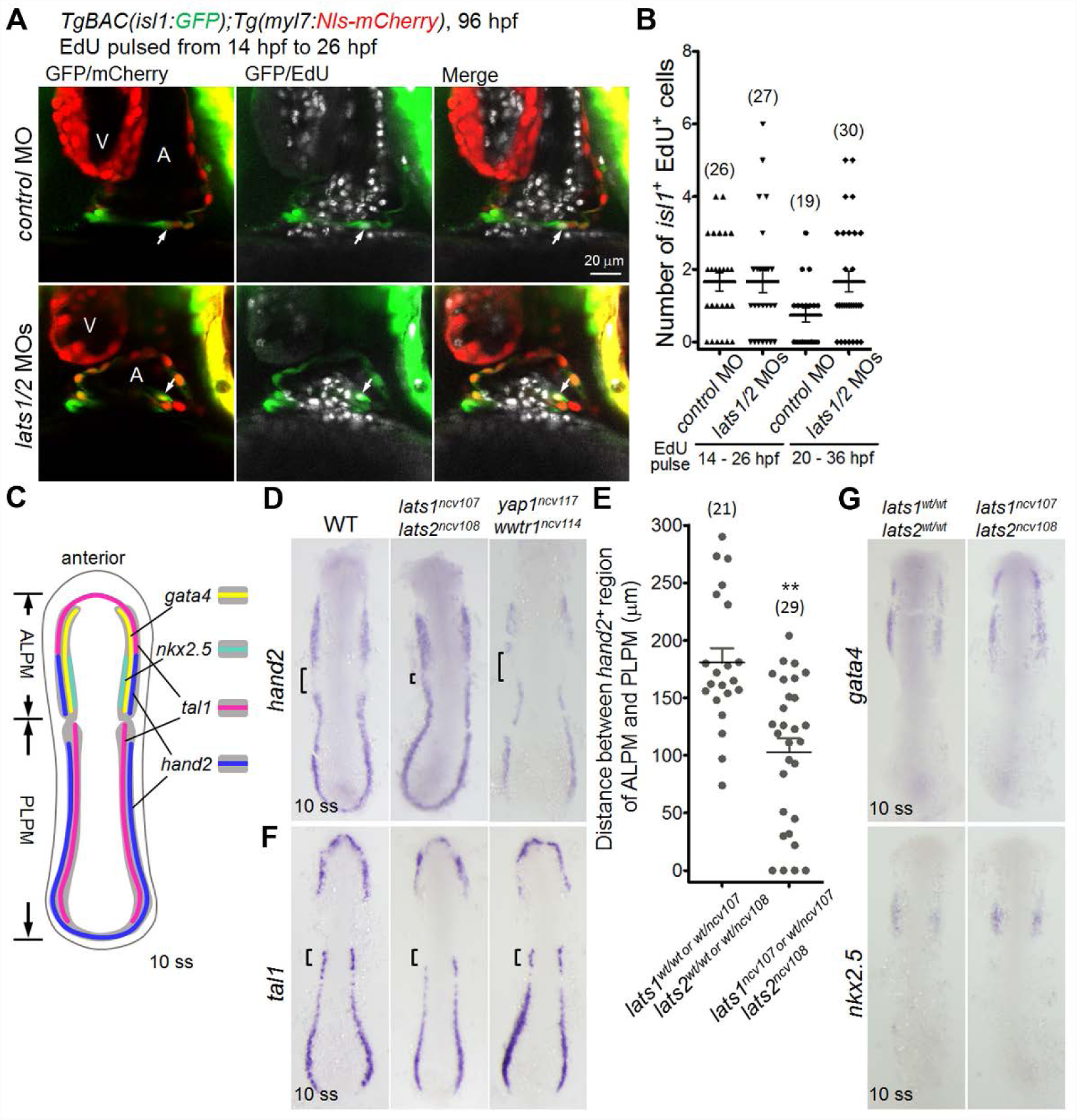
Knockout of *lats1/2* leads to an increase in the expression of *hand2* in the boundary between ALPM and PLPM. **(A)** Single-scan confocal images of the *TgBAC*(*isl1:GFP);Tg(myl7:Nls-mCherry*) embryos injected with the MO indicated at the left and pulsed with EdU from 14 hpf to 26 hpf at 96 hpf. Arrows indicate the EdU-incorporated both *isl1* and *myl7* promoter-activated cells in the IFT of atrium. A, atrium; V, ventricle. Ventral view, anterior to the top. **(B)** The number of EdU-positive *isl1* promoter-activated CMs among EdU-1 positive cells of the embryos treated with the MO as indicated at the bottom. Embryos pulsed with EdU from 14 hpf to 26 hpf (left two columns) and from 20 hpf to 36 hpf (right two columns). **(C)** Schematic illustration of gene expression patterns in the LPM of the wild type (WT) embryos at 10 somite stage (ss). Expression domain of *tal1, gata4, nkx2.5, and hand2* are depicted as magenta, yellow, green, and blue, respectively. Dorsal view, anterior to the top. **(D, F, G)** WISH analyses of the embryos at 10 ss using antisense probe indicated at the left of the panels. **(D, F)** Genotypes are indicated at the top as WT (left panels), *lats1/2* DKO (center panels), and *yap1/wwtr1* DKO (right panels). **(D)** Brackets indicate the gap between *hand2*-positive regions of ALPM and PLPM. **(E)** Quantitative measurement of the distance indicated by the brackets of **(D)** in the either *lats1^wt/wt^ lats2^wt/wt^* or *lats1^wt/ncv107^ lats2^wt/ncv108^* embryos and either *lats1^wt/ncv107^lats2^ncv108^* embryos or *lats1/2* DKO embryos. **(F)** Brackets indicate the *tal1*-positive rostral end of PLPM in the WT. **(G)** Genotypes are indicated at the top as WT (left panels), and *lats1/2* DKO (right panels). Dorsal view, anterior to the top. The single-scan (2 μm) confocal images and ISH images are a set of representative images of at least four independent experiments. ^**^p < 0.01.

We, therefore, asked whether Lats1/2 restrict SHF cell specification from the ALPM. To test this hypothesis, we examined the expression of both ALPM and PLPM genes: *gata4* as a marker for multipotent myocardial-endothelial-myeloid progenitor of ALPM; *nkx2.5* as a marker for ventricular heart field; *tal1* as a marker for hematopoietic cell progenitor; and *hand2* as a marker for both heart field and PLPM at 10 ss (Figure 5C). There was a gap of *hand2* expression between ALPM and PLPM in the WT embryos but the gap length was significantly shorten in either *lats1^wt/ncv107^lats2^ncv108^* embryos or *lats1/2* DKO embryos as well as the *lats1/2* morphants (Figure 5D, E, Figure 5—Figure supplement 1A and Figure 5—source data 1). In clear contrast, *tal1* expression was decreased in the rostral end of PLPM in the *lats1/2* DKO embryos and *lats1/2* morphants (Figure 5F and Figure 5—Figure supplement 1B). Although *hand2* expression was decreased in the both ALPM and PLPM in the *yap1*/*wwtr1* DKO embryos, the expression of *tal1* was unaffected (Figure 5D, F). The expression of *gata4* and *nkx2.5* was unaffected in the both *lats1/2* DKO embryos and *lats1/2* morphants (Figure 5G, and Figure 5—Figure supplement 1C), these results were consistent with the results of qRT-PCR using embryos at 24 hpf (Figure 2A). We further examined other genes regulating heart field; *etv2* as a marker for blood-vessel progenitor (Schoenebeck et al., 2007) and *hoxb5b* as *a* regulatory molecule of RA signaling in the forelimb field (Waxman et al., 2008). The expression of both *etv2* and *hoxb5b* was comparable between the control and the *lats1/2* morphants (Figure 5—Figure supplement 1C). Collectively, these results suggest that Lats1/2 negatively regulates Yap1/Wwtr1-dependent differentiation of LPM to the SHF in the boundary between ALPM and PLPM.

### Yap1/Wwtr1 drive Bmp-Smad signaling essential for SHF formation

Bone morphogenetic proteins (Bmps)-mediated signal affects the various context of heart development via Smad phosphorylation-dependent transcriptional activation. Bmp-Smads signaling is known to be essential for SHF formation, FHF-derived CM development, endocardium development, and epicardium development (Prall et al., 2007; Schlueter et al., 2006; Tirosh-Finkel et al., 2010; Yang et al., 2006). Yap1 promotes *Bmp2b* expression in the neocortical astrocyte differentiation (Huang et al., 2016). In the zebrafish embryos, *bmp2b*, but not *bmp4*, is expressed in the LPM (Chung et al., 2008). We hypothesized that Yap1/Wwtr1 are involved in *bmp2b*-dependent signal during early cardiogenesis. To investigate whether Bmp-Smad signaling is activated in the ALPM, we examined the Bmp-dependent transcription using Tg fish in which Bmp responsive element (BRE) drives GFP expression; *Tg(BRE:GFP)* (Collery and Link, 2011). At 14 hpf, BRE-positive cells were found in the ALPM (Figure 6A). BRE-positive cells in the caudal end of ALPM moved toward the venous pole (Video 4). At 10 ss, *bmp2b* expression was increased in the ALPM in the *lats1/2* DKO embryos and was decreased in the *yap1/wwtr1* DKO embryos (Figure 6B). Consistently, *bmp2b* mRNAs were increased in the *lats1/2* morphants at 10 ss (Figure 6—Figure supplement 1A). Although we could not detect *bmp4* in the ALPM in the early ss (data not shown), *bmp4* mRNAs were increased in the venous pole of the *lats1/2* morphants at 26 hpf (Figure 6—Figure supplement 1B).

**Figure 6.**
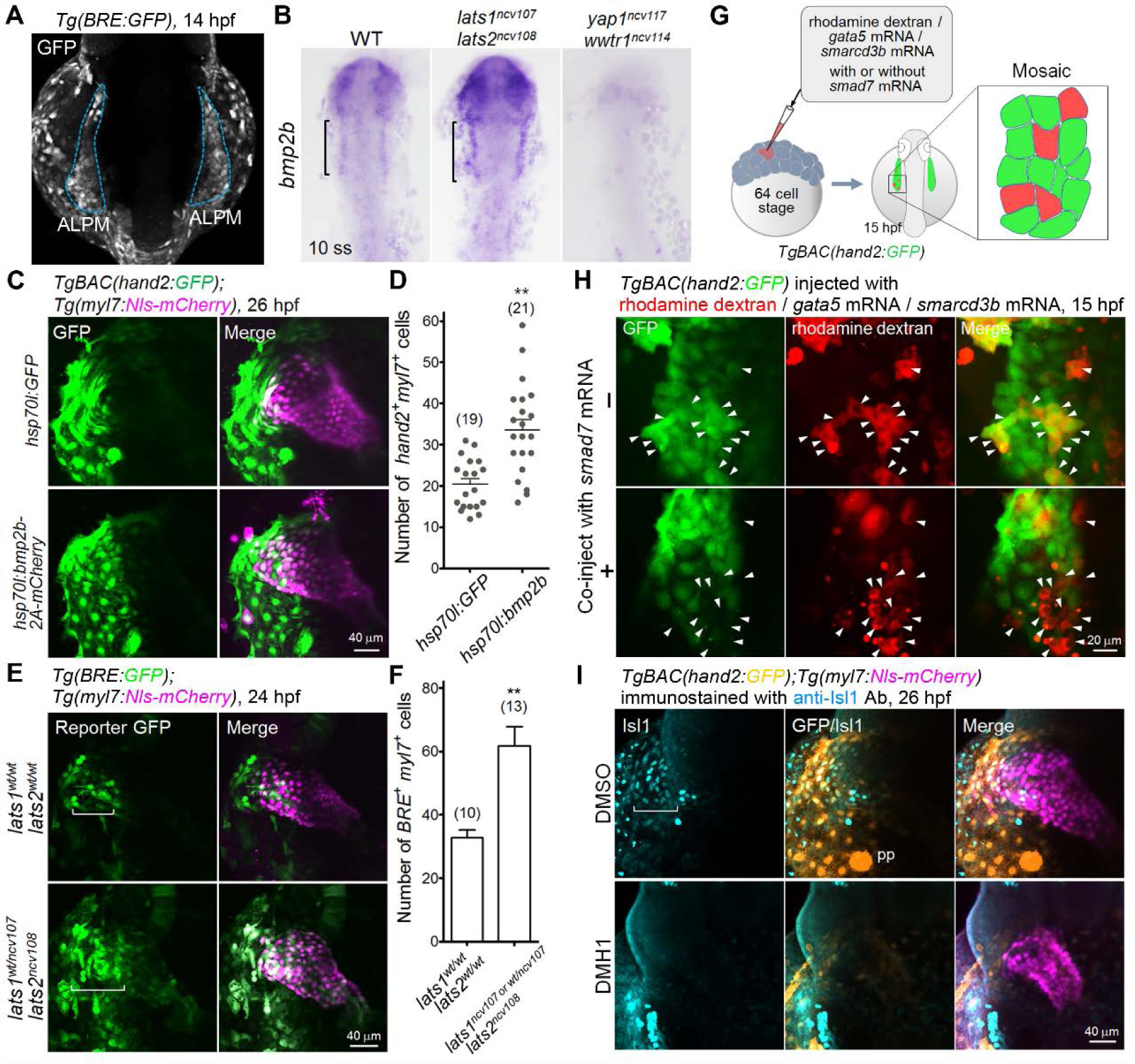
Hippo signaling functions upstream of Bmp-Smad signal that is necessary for Isl1-positive SHF formation. **(A)** 3D confocal stack images of the *Tg(BRE:GFP)* embryos at 14 hpf. Blue broken lines indicate the GFP-positive ALPM. **(B)** WISH analyses of the embryos at 10 ss of the WT, *lats1/2* DKO, and *yap1/wwtr1* DKO indicated at the top using antisense probe for *bmp2b*. Brackets indicate the *bmp2b*-positive ALPM. **(C)** 3D confocal stack images of the *TgBAC(hand2:GFP);Tg(myl7:Nls-mCherry)* embryos injected with pTol2-*hsp70l*:GFP and pTol2-*hsp70l*:bmp2b-2A-mCherry and treated by heat shock at 2 ss for 1 h at 26 hpf. **(D)** Quantitative analyses of the number of both *hand2* and *myl7* promoter-activated cells in the venous pole. Note that overexpression of bmp2b leads to an increase in the number of the *hand2* promoter-activated cardiomyocytes in the venous pole. **(E)** 3D confocal stack images of the *Tg(BRE:GFP);Tg(myl7:Nls-mCherry*) embryos with *lats1^wt/wt^lats2^wt/wt^* allele and *lats1^wt/ncv107^lats2^ncv108^* allele at 24 hpf. Brackets indicate the GFP-positive *myl7* promoter-activated cells in the venous pole. Note that GFP-positive *myl7* promoter-activated cells are increased in the venous pole. **(F)** Quantitative analyses of the number of the both *BRE*-activated GFP-positive and *myl7* promoter-activated mCherry-positive cells in the *lats1^wt/wt^lats2^wt/wt^* embryos and either *lats1^wt/ncv107^lats2^ncv108^* embryos or *lats1/2* DKO embryos. **(G)** Schematic illustration of mosaic analysis. CPC-fated cells by the injection of *gata5* and *smarcd3b* mRNAs were simultaneously injected with *smad7* mRNA and rhodamine dextran into one blastomere at 64 cell stage of *TgBAC(hand2:GFP)* embryos. At 15 hpf, the caudal region of the left ALPM was imaged by confocal microscopy. **(H)** 3D confocal stack images of the *TgBAC(hand2:GFP)* embryos injected with rhodamine dextran, *gata5* mRNA, *smarcd3b* mRNA without *smad7* mRNA (upper panels) and with *smad7* mRNA (bottom panels) at 15 hpf. Arrowheads indicate the rhodamine dextran-labelled cells in the caudal region of the left ALPM. Note that *hand2* promoter-activated GFP signal was suppressed in the cells that express *smad7* mRNA. **(I)** 3D confocal stack images of the *TgBAC*(*hand2:GFP);Tg(myl7:Nls-mCherry*) embryos treated with DMSO (upper panels) or DMH1 (10 μM, bottom panels) from 14 hpf to 26 hpf and immunostained with anti-Isl1 Ab at 26 hpf. Bracket indicates Isl1-positive cells in the venous pole. All images in Figure 6 are dorsal view, anterior to the top. Note that both *hand2* and *myl7* promoter-activated and Isl1-positive cells in the venous pole are absent in the embryos treated with DMH1. The confocal 3D-stack images and ISH images are a set of representative images of at least four independent experiments. ^**^p < 0.01.

The zebrafish bmp2b is essential for dorsoventral patterning before the formation of heart (Kishimoto et al., 1997). To examine the role of bmp2b on the ALPM cells that become the *hand2* promoter-activated cells in the venous pole, we applied the heat shock-dependent overexpression of bmp2b. Overexpression of bmp2b led to an increase in the number of both *hand2* and *myl7* promoter-activated cells in the venous pole of the embryos at 26 hpf (Figure 6C, D and Figure 6—source data 1). These data suggest that bmp2b-9 dependent signal can promote *hand2* expression when ALPM cells become the cells in the venous pole.

The number of Bmp signal-activated cells marked by GFP in the venous pole was increased in either *lats1^wt/ncv107^lats2^ncv108^* embryos or *lats1/2* DKO embryos as well as the *lats1/2* morphants at 24 hpf (Figure 6E, F, Figure 6— Figure supplement 2A and Figure 6—source data 1). Because Bmps induce phosphorylation of Smad1/5/9 (Smad9 is also known as Smad8) (Heldin et al., 1997), we examined phosphorylation of Smad1/5/9 in the venous pole at 26 hpf. The number of phosphorylated Smad1/5/9-positive and *hand2* promoter-activated cells was increased in the venous pole of the *lats1/2* morphants at 26 hpf (Figure 6—Figure supplement 2B). These results suggest that Yap1/Wwtr1 promote *bmp2b* expression and induce the subsequent signaling and that Lats1/2 restrict this Yap1/Wwtr1-dependent signaling to form the proper venous pole.

To investigate whether Bmp-Smad signal functions to promote *hand2* expression in a cell-autonomous manner, we performed mosaic analysis. Smad7, an inhibitory-Smad, blocks the Bmp-Smad signal by interacting with activated Bmp type I receptors and thereby preventing the activation of receptor-regulated Smads (Souchelnytskyi et al., 1998). The number of *isl1* promoter-activated cells was decreased in the *TgBAC(isl1:GFP);Tg(myl7:Nls-*9 *mCherry)* embryos injected with *smad7* mRNA at 26 hpf (Figure 6—Figure supplement 2C). We then tested cell autonomous function by injection of *smad7* mRNA in CPC-fated cells (Fukui et al., 2014; Lou et al., 2011) to see *hand2* expression in the caudal region of the left ALPM (Figure 6G). *hand2* promoter-3 activated GFP signal was suppressed in the cells injected with *smad7* mRNA together *gata5* and *smarcd3b* mRNAs, suggesting the cell-autonomous regulation (Figure 6H).

Finally, to confirm the necessity of Bmp-Smad-regulated signal during the SHF formation, we treated *TgBAC(hand2:GFP);Tg(myl7:Nls-mCherry)* embryos with a Bmp inhibitor, DMH1, from 14 hpf to 26 hpf. The efficiency of DMH1 was confirmed by decreased phosphorylation of Smad1/5/9 (Figure 6—Figure supplement 2B). The expression of Isl1 and the promoter activity of *hand2* were greatly reduced in the embryos treated with DMH1, although the activity of *myl7* promoter monitored by mCherry expression was not affected in the FHF-derived region (Figure 6I). These data suggest that Lats1/2 restrict Yap1/Wwtr1-promoted Bmp2b-dependent signaling cell-autonomously required for both *hand2* and *isl1* promoter-activated SHF formation.

## Discussion

We here for the first time show that Hippo signal is involved in the determination of cell fate of LPM cells (Figure 7). While Yap1/Wwtr1-promoted signal increased the domain of SHF, Lat1/2 restricted it, because Lat1/2 likely to induce export of Yap1/Wwtr1 from the nucleus. The formation of the boundary between ALPM and PLPM as well as whole LPM was under the regulation of Hippo signaling. A cooperative mechanisms by the several signaling pathways might be involved in the definition of heart field. In the forelimb field, *hoxb5b* represses the extension of posterior end of heart field that differentiate into atrial but not ventricular CMs (Waxman et al., 2008). Furthermore, the pronephric field in the intermediate mesoderm and the angiogenic field in the rostral region of PLPM are closely associated with restriction of their cell fate in these boundary (Kimmel et al., 1990; Mudumana et al., 2008). The increased number of SHF cells in the *lats1/2* DKO embryos may be attributable to the change of fate determination from *hand2*-negative cells to-positive cells in the boundary between ALPM and PLPM. Indeed, we found that expression of the marker of blood-cell progenitor *tal1* was repressed at the rostral region of PLPM in the *lats1/2* DKO embryos. Although mutants of *lats1/2* exhibited a subtle increase in the number of Isl1-positive atrial SHF cells with no apparent defect of other organs, Hippo signaling is involved in the lineage specification of LPM cells.

**Figure 7.**
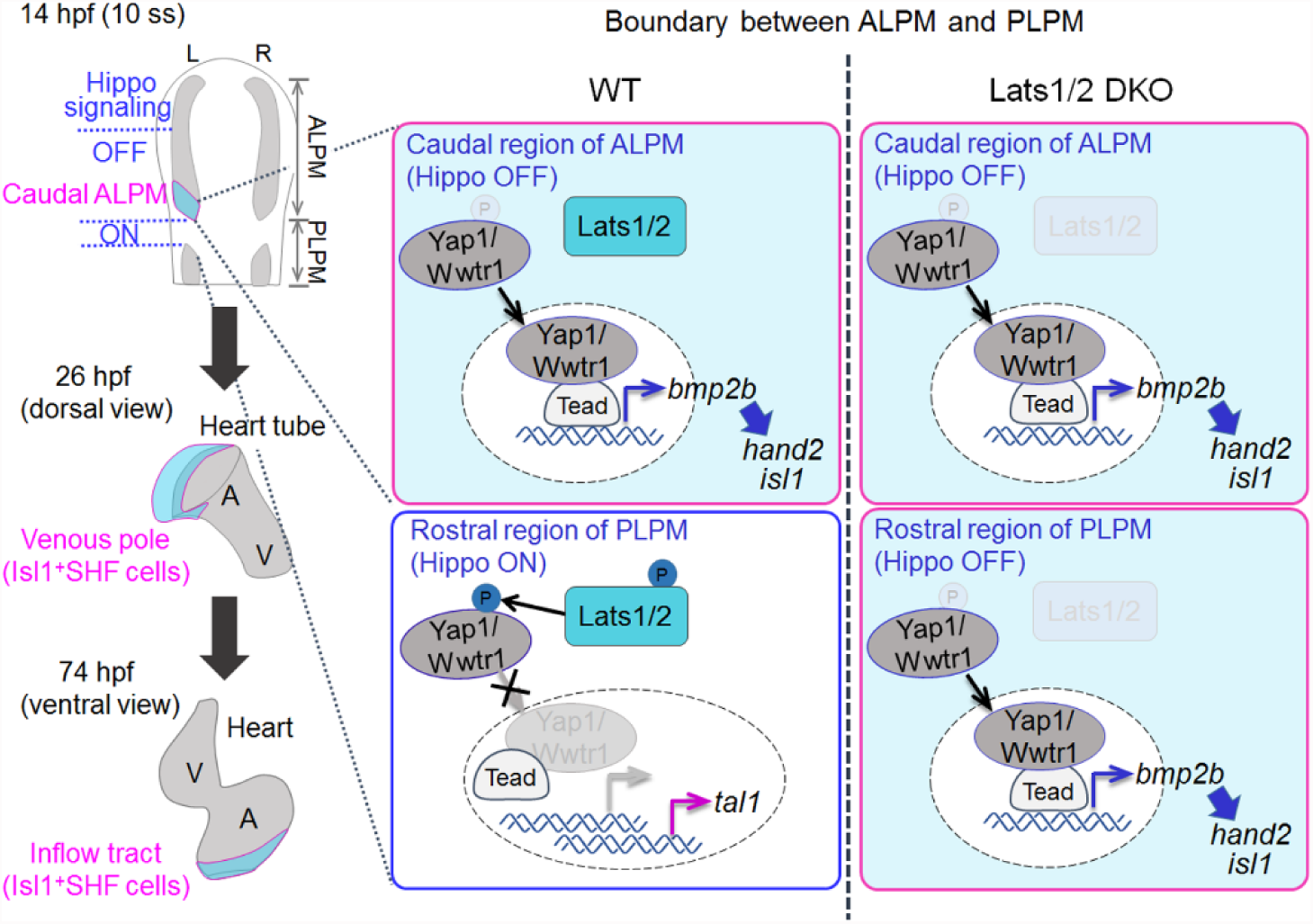
A schematic representation of inflow tract atrial CMs development from the caudal region of the left ALPM. Our study suggest that Lats1/2-Yap1/Wwtr1 signaling controls *bmp2b* expression in the ALPM. Secreted Bmp2b activates Smad signaling that cell autonomously induces the expression of *hand2* to differentiate Isl1 positive SHF cells in the caudal region of the left ALPM. In the *lats1/2* DKO embryos, Yap1/Wwtr1 facilitate Tead-dependent *bmp2b* expression, and thereby expands the expression of *hand2* caudally at the expense of *tal1* expression in the rostral region of PLPM. Consequently, hippo signaling restricts the number of the venous pole cells and lately inflow tract atrial CMs by regulating Bmp-Smad signaling in the boundary between ALPM and PLPM.

Zebrafish Isl1-positive SHF cells might correspond to the mammalian posterior-SHF. In the mouse embryo, the anterior and posterior SHF differentiate into OFT/right ventricular myocardium and IFT/atrial myocardium, respectively (Galli et al., 2008; Verzi et al., 2005). The posterior-SHF in the heart field is located caudally (Galli et al., 2008; Abu-Issa and Kirby, 2008). Pitx2, a member of the paired-like family of homeodomain transcription factors, is expressed in the left LPM for embryonic left-right asymmetry and determines the posterior-SHF formation (Galli et al., 2008). Tead reporter activation occurred in the left side of the caudal zone of ALPM that becomes Isl1-positive venous pole atrial CMs but not ventricular CMs. Caudally-located left and right side of heart fields constituted the venous pole and the arterial pole of cardiac tube, respectively. In addition, mammalian SHF cells have multi-potential to differentiate into endocardium and smooth muscle cells in addition to myocardium (Chen et al., 2009b). By generating BAC transgenic fish, we found that zebrafish *isl1* promoter-activated SHF cells gave rise to atrial myocardium, endocardium, and epicardium, except for ventricular myocardium. Combining with our results and previous reports, the properties of zebrafish Isl1-positive SHF cells are similar to that of the mammalian posterior-SHF cells.

Lats1/2-Yap1/Wwtr1-Tead signaling functions upstream of Bmp2b-Smad activation in the ALPM that is necessary for the formation of *hand2* and *isl1* promoter-activated cells. Although the previous reports have shown that Isl1-positive and Mef2-positive cells reside in the venous pole (de Pater et al., 2009; Hinits et al., 2012), the molecular mechanism underlying how these cells give rise to the CPCs in the venous pole has remained unclear. To date, extracellular stimuli TGFβ, FGF, and BMP have been reported to regulate arterial pole formation in zebrafish (de Pater et al., 2009; Hami et al., 2011; Zhou et al., 2011). In addition, transcription factors including Tbx1, Mef2c, and Nkx2.5 are known to control the development of arterial but not venous pole formation (Guner-Ataman et al., 2013; Hinits et al., 2012; Lazic and Scott, 2011). We revealed that Yap1/Wwtr1-Tead dependent transcription is required for *isl1*-promoter activated SHF formation, because the forced expression of Yap1/Wwtr1-Tead binding dominant-negative form (ytip-mCherry) suppressed the formation of *isl1* promoter-activated SHF cells. Furthermore, Bmp reporter activity was observed in the ALPM. By analyzing the Lats1/2 mutants and Yap1/Wwtr1 mutants, we demonstrates that Hippo signal controls *bmp2b* expression in the ALPM. Bmp-Smad inhibition expands *tal1* expression domain to restrict LPM fate (Gupta et al., 2006). In addition, *hand2* expression is diminished in the mutant of *alk3*, a Bmp type I receptor 1a, at 12 ss (de Pater et al., 2012). In our hands, Bmp-Smad inhibition resulted in cell-autonomous suppression of the *hand2* promoter-positive GFP expression at 15 hpf. Therefore, Bmp2b expression positively regulated by Yap1/Wwtr1 balances the cell fate to the heart field and to the blood cells in the boundary between ALPM and PLPM.

In summary, we demonstrate that Yap1/Wwtr1-Tead signal promotes Bmp2b expression and activates subsequent Smad signaling in the cells located in the left side of caudally-located ALPM (Figure 7). This signaling determines the Isl1-positive SHF cells in the venous pole that specifically become the IFT atrial CMs lately.

## Materials and methods

### Zebrafish (*Danio rerio*) strain, transgenic lines and mutant lines

The experiments using zebrafish were approved by the institutional animal committee of National Cerebral and Cardiovascular Center and performed according to the guidelines of the Institute. We used the AB strain as wild-type. The following zebrafish transgenic lines were used for experiments: *Tg(eef1a1l1:galdb-hTEAD2*Δ*N-2A-mCherry)* fish (Fukui et al., 2014), *Tg(myl7:Nls-mCherry)* fish (Fukui et al., 2014), *TgBAC*(*hand2:GFP*) fish (Yin et al., 2010), *Tg(BRE:GFP)* fish (Collery and Link, 2011), and *Tg(uas:GFP)* fish (Asakawa et al., 2008). The *Tg(myl7:galdb-hTEAD2*Δ*N-2A-mCherry)* fish, *Tg(myh6:Nls-tdEosFP)* fish, and *TgBAC(isl1:GFP)* fish were generated as described supplementary experimental procedures. The knockout alleles as *ncv107* for *lats1*, *ncv108* for *lats2*, and *ncv117* for *yap1* genes were generated by TALEN techniques as described supplementary experimental procedures. The *ncv114* allele for *wwtr1* was previously reported (Nakajima et al., 2017).

### Image acquisition by microscopies and image processing

To clearly obtain the images of embryos, pigmentation of embryos was suppressed by addition of 1-phenyl-2-thiourea (PTU) (Sigma-Aldrich, St. Loiuis, MO) into breeding E3 media. Embryos were dechorionated and mounted in 1% low-melting agarose dissolved in E3 medium. Confocal images were taken with a FV1200 confocal microscope system (Olympus, Tokyo, Japan) equipped with water immersion 20x lens (XLUMPlanFL, 1.0 NA, Olympus). Images were processed with FV10-ASW 4.2 viewer (Olympus). Distance between *hand2* positive-region of ALPM and PLPM was measured by a DP2-BSW software (Olympus). Cell tracking data containing nuclei positions and instantaneous velocities were analyzed by Imaris8.4.1 software (Bitplane, Zurich, Switzerland).

### Generation of knockout zebrafish by TALEN

To make knockout zebrafish, we used transcription activator-like effector nuclease (TALEN) Targeter 2.0 (https://tale-nt.cac.cornell.edu) to design TALEN pair that targets *lats1*, *lats2* and *yap1*. The target sequence of TAL-*lats1*, TAL-8 *lats2, and* TAL-*yap1* were 5′-TCAGCAAATGCTGCAGGAGATccgagagagcctgcgaAACCTCTCCCCGTCCTCC AA-3′, 5′-TCTCGAGGAGAGGGTGgtcgaggtggagactCAAAGGGCAAAGACCA-3′, and 5′-CCGAACCAGCACAACCctccagccggccaccagaTCGTCCATGTTCGGGG-3′,respectively (capital letters were sequences of left [TAL-*lats1*-F, *lats2*-F, and *yap1*-F] and right [TAL-*lats1*-R, *lats2*-R, and *yap1*-R] arms, respectively). These expression plasmids of the TALEN-pair were constructed by pT3TS-6 GoldyTALEN. TALEN mRNAs were synthesized in vitro by T3 mMessage mMACHINE kit (Thermo Fisher Scientific, Waltham, MA). To induce double strand breaks in the target sequence, both 50 pg of TAL-*lats1-F* / -*lats1-R* mRNAs, TAL-*lats2-F* / -*lats2-R* mRNAs, and TAL-*yap1-F* / -*yap1-R* mRNAs were injected into 1-2-cell stage Tg embryos, respectively. Each injected founder (F0) fish was outcrossed with wild-type fish to obtain F1 progeny from the individual founders. Generation of *wwtr1* knockout zebrafish was previously reported (Nakajima et al., 2017). To analyze TALEN induced mutations, genomic DNA from F1 embryos was lysed by 50 μl of NaOH solution (50 mM) at 95°C for 5 min, and added 5 μl of Tris-HCl (pH8.0, 1.0 M) on ice for 10 min. After centrifugation (13,500 rpm, 5 min), PCR reaction was performed by KOD FX Neo DNA polymerase (TOYOBO, Osaka, Japan). The genotyping PCR primers were used for amplification: *lats1* (5′-GGCACTTAACATATGCTTTTACATG-3’ and 5′-TTTGCTGCTGTCTGCGGAGCTGTT-3′); *lats2* (5′-AGAGTTTGTGTGAGAGAAAACAGG-3’ and 5′-GCATTGACCAGATCCTGTAGCATC-3′); *yap1* (5′-TCCTTCGCAAGGCTTGGATAATTG-3’ and 5′-TTGTCTGGAGTGGGACTTTGGCTC-3′); *wwtr1* (5′-GGACGAAAAACAGGAAAAGTTC-3’ and 5′-ACTGCGGCATATCCTTGTTC-3′). These amplified PCR products were analyzed using MCE-202 MultiNA microchip electrophoresis system (SHIMADZU, Kyoto, Japan) with the DNA500 reagent kit (SHIMADZU).

### Microinjection of oligonucleotide and mRNA

We injected 200 pg *ytip-mCherry* mRNA (Fukui et al., 2014), 100 pg zebrafish-*smad7* mRNA, 1.2 ng *lats1-atg* MO (5′-CCTCGGGTTTCTCGGCCCTCCTCAT-3′) (Chen et al., 2009a), 1.2 ng *lats2-atg* MO (5′-CATGAGTGAACTTGGCCTGTTTTCT-3′) (Chen et al., 2009a), 3 ng *pkd2-atg* MO (5′-ACTGGAGTTCATCGTGTATTTCTAC-3′) (Bisgrove et al., 2005), 8 ng *ajuba-atg* MO (5′-TGAGTTTGATGCCAAGTCGATCCAT-3′) (Witzel et al., 2012), and 5 ng *control* MO (5′-CCTCTTACCTCAGTTACAATTTATA-3′) as previously reported (Fukui et al., 2014). These morpholinos were purchased from Gene Tools (Philomath, OR). Capped Messenger RNAs were synthesized using SP6 mMessage mMachine system (Thermo Fisher Scientific). Microinjection was performed by using FemtoJet (Eppendorf, Hamburg, Germany). MOs, mRNA, and Tol2 plasmids were injected into one-cell to two-cell stage blastomere.

### Heat shock treatment

*TgBAC(hand2:GFP);Tg(myl7:Nls-mCherry)* embryos were injected with 25 pg pTol2-*hsp70l*:GFP or 25 pg pTol2-*hsp70l*:bmp2b-2A-mCherry plasmids along with 50 pg *tol2 transposase* mRNA and heat shocked at 2 ss for 1 hr at 39°C.

### Mosaic assay

Mosaic assay was performed as previously described (Fukui et al., 2014). For injection of *smad7* mRNA into CPC-fated cells, *smad7* mRNA (4 pg) was co-2 injected with *gata5* mRNA (1.5 pg) and *smarcd3b* mRNA (1.5 pg) together with rhodamine dextran (70,000 MW, lysine fixable [Thermo Fisher Scientific]) as a cell tracer into one blastomere at the 64 cell stage of *TgBAC(hand2:GFP)* embryos. The *hand2* promoter-activated caudal region of the left ALPM of the embryos was confocal imaged at 15 hpf (12 ss).

### EdU incorporation assay

*TgBAC(isl1:GFP);Tg(myl7:Nls-mCherry)* embryos injected with *control* MO or *lats1/2* MOs were incubated with 2 mM of 5-Ethynyl-2-deoxyuridine (EdU) from 14 to 26 hpf or 20 to 36 hpf, and subsequently fixed by 4% PFA at 96 hpf. EdU incorporated cells were labelled by Click-iT EdU Alexa Fluor 647 Imaging Kits (Thermo Fisher Scientific) following manufacturer’s instructions. Images were taken by FV1200 confocal microscope system. The number of EdU-positive *isl1* promoter-activated CMs were measured by the overlapping cell of Alexa Fluor 647-positive signal and *isl1* and *myl7* promoter-activated signals.

### Whole-mount in situ hybridization (WISH)

The antisense *hand2, bmp2b, bmp4, gata4, nkx2.5, etv2, tal1, and hoxb5b* RNA probes labeled with digoxigenin (DIG) were prepared by using an RNA labeling kit (Roche, Basel, Switzerland). WISH was performed as previously described (Fukui et al., 2014). Colorimetric reaction was carried out using BM purple (Roche) as the substrate. To stop reaction, embryos were washed by PBS-T and fixed by 4% PFA for 20 min at room temperature and subsequently substituted by glycerol. Images were taken by SZX-16 Stereo Microscope (Olympus).

### Immunohistochemistry

Embryos at 26 hpf were fixed by MEMFA (3.7% formaldehyde, 0.1 M MOPS, 2 mM EGTA, 1 mM MgSO4) for 2 hr at room temperature. After fixation, the solution was changed to 50% Methanol / MEMFA for 10 min, changed to 100% Methanol at room temperature, and stored in 100% Methanol at -30°C overnight. After rehydration, embryos were washed three-times for 10 min in PBBT (PBS with 2 mg/mL BSA and 0.1% TritonX-100). Embryos were blocked in PBBT with 10% goat serum for 60 min at room temperature, and subsequently incubated overnight at 4°C with primary antibodies, 1:300 diluted chicken anti-GFP antibody (ab13970, Abcam, Cambridge, UK), 1:300 diluted mouse anti-mCherry antibody (632543, Clontech, Mountain View, CA), and 1:100 diluted rabbit anti-Islet1 antibody (GTX128201, Genetex, Irvine, CA) or 1:100 diluted rabbit anti-pSmad1/5/9 antibody (13820S, Cell Signaling TECHNOLOGY, Danvers, MA) in blocking solution. Embryos were washed with PBBT for five-times over the course of 2 hours, with blocking solution for 60 min at room temperature, and incubated overnight at 4°C with secondary antibodies, anti-chicken Alexa Fluor 488 IgG (A-11039, Thermo Fisher Scientific), anti-5 mouse Alexa Fluor 546 IgG (A-11030, Thermo Fisher Scientific), and anti-rabbit Alexa Fluor 633 IgG (A-21070, Thermo Fisher Scientific) diluted 1:300 in blocking solution. Embryos were washed with PBBT for five-times over the course of 2 hours, and stored in PBS at 4°C prior to imaging.

### Quantitative real time PCR (q-PCR)

Total RNAs were collected from whole-embryonic cells by using TRizol (Thermo Fisher Scientific) following the manufacturer’s instructions. For q-PCR, reverse transcription and RT-PCR were performed with QuantiFast SYBR Green PCR kit (Qiagen, Hilden, Germany) in Mastercycler Realplex (Eppendorf). The following primer set were used for amplification: *nkx2.5*-S (5′-GCTTTTACGCGAAGAACTTCC-3′), *nkx2.5*-AS (5′-GATCTTCACCTGTGTGGAGG-3′); *gata4*-S (5′-AAGGTCATCCCGGTAAGCTC-3′), *gata4*-AS (5′-TGTCACGTACACCGGAGAAG-3′); *hand2*-S (5′-TACCATGGCACCTTCGTACA-3′), *hand2*-AS (5′-CCTTTCTTCTTTGGCGTCTG-3′); *eef1a1l1*-S (5′-CTGGAGGCCAGCTCAAACAT-3′), *eef1a1l1*-AS (5′-ATCAAGAAGAGTAGTACCGCTAGCATTAC-3′) (Fukui et al., 2014).

### Plasmids

cDNA fragments encoding zebrafish Hand2, Bmp2b, Bmp4, Gata4, Nkx2.5, Etv2, Tal1, Hoxb5b and Smad7 were amplified by PCR using a cDNAs library derived from zebrafish embryos and subcloned into pCR4 Blunt TOPO vector (Thermo Fisher Scientific). The following primer set were used for amplification: *hand2*-S (5′-CGGGATCCCGCCATGAGTTTAGTTGGAGGGTT-3’ [containing BamHI sequence]), *hand2*-AS (5′-GCTTTAGTCTCATTGCTTCAGTTCC-3′); *bmp2b*-S (5′-ATGTCGACACCATGGTCGCCGTGGTCCGCGCTCTC-3’ [containing SalI sequence]), *bmp2b*-AS (5′-TCATCGGCACCCACAGCCCTCCACC-3′); *bmp4*-S (5′-CGGGATCCCATGATTCCTGGTAATCGAATGC-3’ [containing BamHI sequence]), *bmp4*-AS (5′-CATTTGTACAACCTCCACAGCAAG-3′); *gata4*-S (5′-GTGAATTCATGTATCAAGGTGTAACGATGGCC-3’ [containing EcoRI sequence]), *gata4*-AS (5′-GAGCTTCATGTAGAGTCCACATGC-3′); *nkx2.5*-S (5′-GCTCTAGATTCCATGGCAATGTTCTCTAGCCAA-3’ [containing XbaI sequence]), *nkx2.5*-AS (5′-GATGAATGCTGTCGGTAAATGTAG-3′); *etv2*-S (5′-GTGAATTCCTGGATTTTACACAGAAGACTTCAGA-3’ [containing EcoRI sequence]); *etv2*-AS (5′-CCACGACTGAGCTTCTCATAGTTC-3′); *tal1*-S (5′-GTGAATTCGAAATCCGAGCAATTTCCGCTGAG-3’ [containing EcoRI sequence]), *tal1*-AS (5′-CTTAGCATCTCCTGAAGGAGGTCGT-3′); *hoxb5b*-S (5′-GTGAATTCCCAAATGAGCTCTTATTTTCTAAACTCG-3’ [containing EcoRI sequence]), *hoxb5b*-AS (5′-GATGTGATTTGATCAATTTTGAAACGCGC-3′); *smad7*-S (5′-AGGGATCCTCCCGCATGTTCAGGACCAAACGAT-3’ [containing BamHI sequence]), *smad7*-AS (5′-GAAGGCCTTTATCGGTTATTAAATATGACCTCTAACC-3’ [containing StuI sequence]). The cDNAs of zYtip, Gata5, and Smarcd3b were previously amplified and cloned into the pCS2 vector (Clontech) (Fukui et al., 2014). The DNA encoding Smad7 was subcloned into the pCS2 vector to construct the pCS2-smad7. pTol2-*hsp70l* was previously reported (Kashiwada et al., 2015). The cDNAs encoding GFP and Bmp2b-2A-mCherry were subcloned into the pTol2-*hsp70l* vector to construct the pTol2-*hsp70l*:GFP and pTol2-*hsp70l*:bmp2b-2A-mCherry. All the cDNAs amplified by PCR using cDNA libraries were sequenced. Mutations were also confirmed by sequencing.

### Generation of Transgenic Lines

To monitor the atrial CM development, we established a transgenic (Tg) zebrafish lines expressing nuclear localization signal (Nls)-tagged tandem Eos fluorescent protein under the control of *myosin heavy chain 6* (*myh6*) promoter; *Tg*(*myh6:Nls-tdEosFP*). pTol2-*myh6* vector was constructed by modifying pTol2 vector and inserting the *myh6* promoter as a driver of expression of the target molecule. The primers to amplify the *myh6* promoter were 5′-AGAGCTAAAGTGGCAGTGTGCCGAT-3’ and 5′-TCCCGAACTCTGCCATTAAAGCATCAC-3′. An oligonucleotide encoding Nls derived from SV40 (PKKKRKV) was inserted into pcDNA-tdEosFP (MoBiTec, Göttingen, Germany) to generate the plasmids expressing Nls-tagged tandem Eos fluorescent protein (Nls-tdEosFP). The Nls-tdEosFP cDNA was subcloned into the pTol2-*myh6* vector to construct the pTol2-*myh6*:Nls-tdEosFP plasmids.

To monitor the CM-specific Yap1/Wwtr1-dependent transcriptional activation, we developed a Tg line which expresses human (h) TEAD2 lacking amino-terminus (1-113 aa) fused with Gal4 DNA binding domain followed by 2A mCherry under the control of *myosin light polypeptide 7* (*myl7*) promoter; *Tg(myl7:galdb-hTEAD2*Δ*N-2A-mCherry).* This Tg fish was crossed with *Tg(uas:GFP)* reporter fish to obtain *Tg(myl7:galdb-hTEAD2*Δ*N-2A-*7 *mCherry);Tg(uas:GFP)*. The pTol2-*myl7* vector and the pcDNA3.1 vector containing human *TEAD2*Δ*N* cDNA fused to the DNA binding domain of Gal4 (pcDNA3.1-galdb-hTEAD2ΔN) were constructed as previously described (Fukui et al., 2014). The Gal4db-hTEAD2ΔN cDNA was subcloned into the pTol2-*myl7* vector to construct the pTol2-*myl7*:galdb-hTEAD2ΔN plasmids. All the cDNAs amplified by PCR using cDNA libraries were confirmed by sequence.

To monitor the SHF development, we established a Tg line which expresses GFP under the control of *isl1* BAC promoter/enhancer; *TgBAC(isl1:GFP).* pRedET plasmid (Gene Bridges, Heidelberg, Germany) was introduced into E. coli containing CH211-219F7 BAC clone encoding *isl1* gene (BacPAC resources) by electroporation (1800V, 25 mF, 200 Ω) to increase the efficiency of homologous recombination, as previously described (Ando et al., 2016). Tol2 long terminal repeats in opposite directions flanking ampicillin resistance cassette were amplified by PCR using Tol2_amp as a template and were inserted into the BAC vector backbone. The cDNA encoding GFP together with a kanamycin resistance cassette (GFP_KanR) was amplified by PCR using pCS2-GFP_KanR plasmid as a template and inserted into the start ATG of the *isl1* gene. Primers to amplify the GFP_KanR for *isl1* gene were 5′-gggccttctgtccggttttaaaagtggacctaacaccgccttactttcttACCATGGTGAGCAAGGGC GAGGAG-3’ and 5′-aaataaacaataaagcttaacttacttttcggtggatcccccatgtctccTCAGAAGAACTCGTCAAG AAGGCG-3’ (small letters; homology arm to BAC vector, and capital letters; primer binding site to the template plasmid). Tol2-mediated zebrafish transgenesis was performed by injecting 30 pg transgene plasmid together with 50 pg *Tol2* mRNA, followed by subsequent screening of F1 founders and establishment of single-insertion transgenic strains through selection in F3 generations.

## Data analysis and statistics

Data were analyzed using GraphPad Prism 7 (GraphPad Software, La Jolla, CA). All columns were indicated as mean ± SEM. Statistical significance of multiple groups was determined by one-way ANOVA with Bonferroni’s post hoc test. The number of atrial and ventricular CMs at 74 hpf were analyzed by Student’s t-test. Statistical significance of two groups was determined by Student’s t-test.

## Acknowledgements

We thank Stainier DY for *TgBAC(hand2:GFP)* fish; Sone M, Babazono T, Hiratomi K, Ueda M, and Toyoshima S, for their technical assistance.

---

The following figure supplements are available for figure 1:

**Figure supplement 1.** Knockout of *lats1/2* genes leads to an activation of the Tead reporter.

**Figure supplement 2.** Tead reporter activation is found in the venous pole CMs of atrium.

**Figure 1—source data 1. Quantification of atrial (Figure 1B) and ventricular (Figure 1C) cardiomyocyte numbers in the embryos with *lats1* and *lats2* mutant.**

The following figure supplements are available for figure 2:

**Figure supplement 1.** Depletion of Lats1/2 results in an increase in the *hand2* promoter-activated cells in the venous pole.

**Figure supplement 2.** *hand2* promoter-activated cells were significantly reduced in the *yap1/wwtr1* double-knockout embryos.

**Figure 2—source data 1. Quantification of the relative mRNAs expression levels (Figure 2A) and the number of both *hand2* and *myl7* promoter-activated cells (Figure 2D).**

The following figure supplement is available for figure 3:

**Figure supplement 1.** Cells in the caudal region of right ALPM of the *pkd2* morphants with situs inversus migrate toward the right-sided venous pole.

**Figure 3—source data 1. Quantification of mean velocities of *hand2* promoter-activated cells and Tead reporter-activated cells in the caudal region of left and right ALPM.**

The following figure supplement is available for figure 4:

**Figure supplement 1.** Depletion of Lats1/2 leads to an increase of the number of Isl1-positive SHF cells in the venous pole.

**Figure 4—source data 1. The number of *isl1* and *myl7* promoter-activated IFT and OFT cells at 96 hpf (Figure 4C) and *isl1* promoter-activated SHF cells of *lat1/2* mutants (Figure 4E) and *lat1/2* morphants (Figure 4F) at 26 hpf.**

The following figure supplement is available for figure 5:

**Figure supplement 1.** Depletion of Lats1/2 leads to an increase in the expression of *hand2*.

**Figure 5—source data 1. Distance between *hand2*-positive regions of ALPM and PLPM of the control and the *lat1/2* mutants at 10 ss.**

The following figure supplements are available for figure 6:

**Figure supplement 1.** Depletion of Lats1/2 leads to an increase in the *bmps* expression.

**Figure supplement 2.** Depletion of Lats1/2 leads to an activation of Bmp-Smad signaling that is necessary for Isl1-positive SHF formation.

**Figure 6—source data 1. The number of both *hand2* and *myl7* promoter-activated cells at 26 hpf (Figure** 6D**) and BRE-positive cardiomyocytes of the lat1/2 mutants at 24 hpf (Figure 6F).**

**Figure 1—Figure supplement 1.**
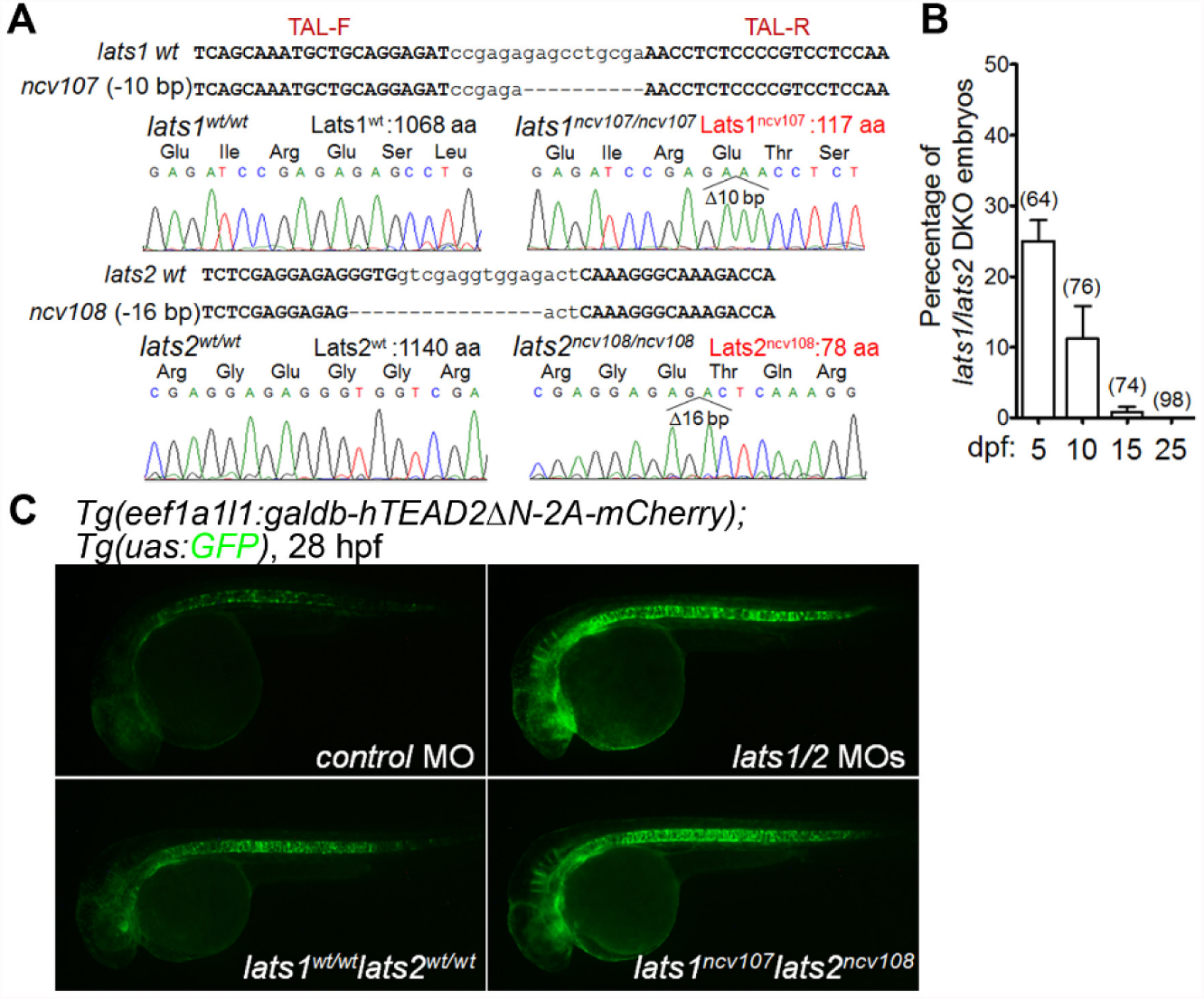
Knockout of *lats1/2* genes leads to an activation of the Tead reporter. **(A)** *lats1* and *lats2* gene mutation by TALEN at the targeted loci. A deletion of ten base pairs in the *ncv107* allele and sixteen base pairs in the *ncv108* allele results in a premature stop codon in exon 3 of *lats1* (resulting mutant Lats1 consists of 117 aa) and exon 3 of *lats2* (resulting mutant Lats2 consists of 78 aa), respectively. Upper and lower case letters denote target and spacer region for the TALEN, respectively. **(B)** The number of double knockout (DKO) embryos by incrossing of *lats1^wt/ncv107^lats2^ncv108^* fishes at the days post-fertilization (dpf) indicated at the bottom. Total number of larvae examined in the experiment is indicated on the top of column. **(C)** Fluorescent images of *Tg(eef1a1l1:galdb-hTEAD2*Δ*N-2A-*7 *mCherry);Tg(uas:GFP*) embryos injected with *control* MO and *lats1/2* MOs (upper panels), and with *lats1^wt/wt^lats2^wt/wt^* allele and *lats1^ncv107^lats2^ncv108^* allele (bottom panels) at 28 hpf. Lateral view, anterior to the left. The fluorescent images **(C)** are a set of representative images of four independent experiments.

**Figure 1—Figure supplement 2.**
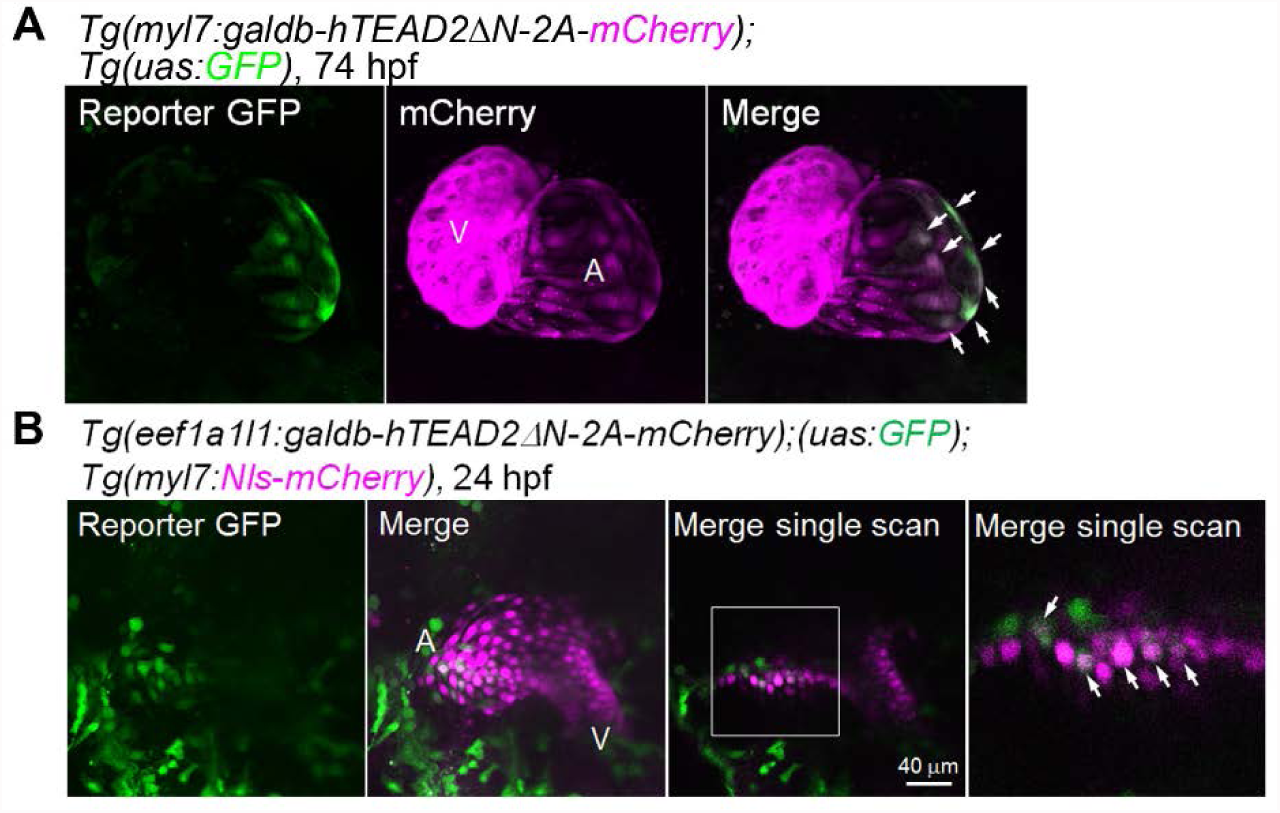
Tead reporter activation is found in the venous pole CMs of atrium. **(A)** 3D confocal stack images of *Tg*(*myl7:galdb-hTEAD2*Δ*N-2A-mCherry);Tg(uas:GFP*) embryos at 74 hpf. Arrows indicate the TEAD reporter GFP-positive atrial CMs. Reporter GFP image (left), mCherry image (center), and merged image of GFP and mCherry (right). A, atrium; V, ventricle. Ventral view, anterior to the top. **(B)** Confocal images of *Tg(eef1a1l1:galdb-*9 *hTEAD2*Δ*N-2A-mCherry);Tg(uas:GFP*);*Tg(myl7:Nls-mCherry)* embryos at 24 hpf. The boxed region is enlarged and shown in the most right panel. Arrows indicate the Tead reporter-positive cells that might differentiate into atrial CMs. Regions in the heart tube that would give rise to atrium (A) and ventricle (V) are marked. 3D-stack images (left two panels) and single scan images (right two panels). Dorsal view, anterior to the top. The confocal 3D-stack images are a set of representative images of at least four independent experiments.

**Figure 2—Figure supplement 1.**
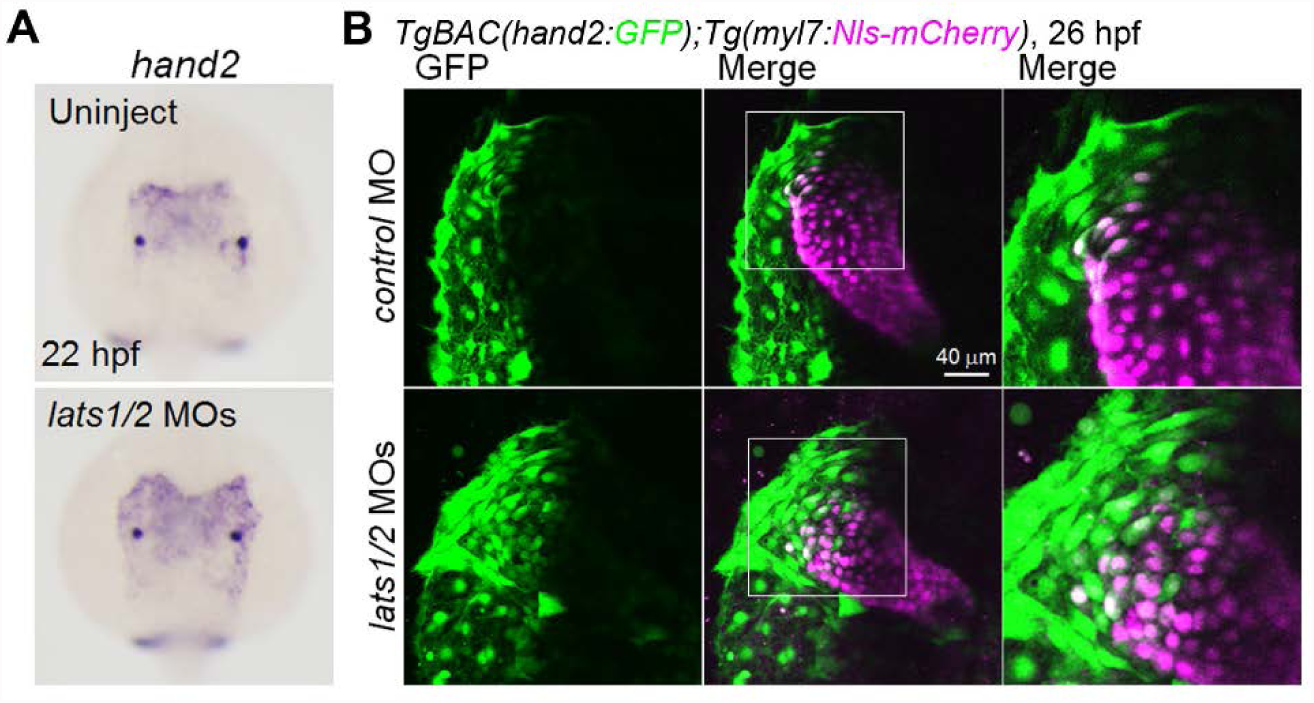
Depletion of Lats1/2 results in an increase in the *hand2* promoter-activated cells in the venous pole. **(A)** Whole mount in situ hybridization (WISH) analyses of the embryos at 22 hpf of the control (uninject) and injected with the *lats1/2* MOs indicated at the top using antisense probe for *hand2*. **(B)** 3D confocal stack images of *TgBAC*(*hand2:GFP);Tg(myl7:Nls-mCherry*) embryos injected with *control* MO and *lats1/2* MOs at 26 hpf. GFP images (left), merged images of GFP image and mCherry image (center), and enlarged images of boxed regions in the center panels (right). Dorsal view, anterior to the top. Images are a set of representative images of eight independent experiments.

**Figure 2—Figure supplement 2.**
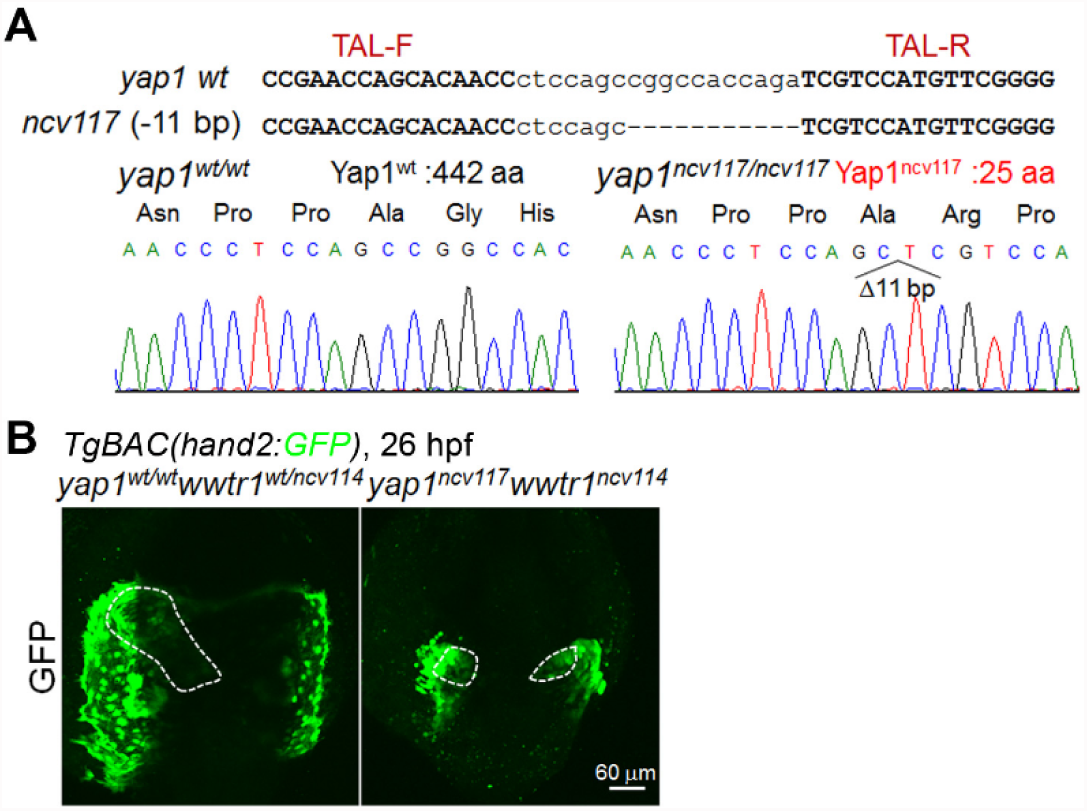
*hand2* promoter-activated cells were significantly reduced in the *yap1/wwtr1* double-knockout embryos. **(A)** *yap1* gene mutation by TALEN at the targeted loci. A deletion of eleven base pairs in the *ncv117* allele results in a premature stop codon in exon 1 of *yap1* (resulting mutant Yap1 consists of 25 aa). Upper and lower case letters denote target and spacer regions for the TALEN, respectively. **(B)** 3D confocal stack images of *TgBAC*(*hand2:GFP)* embryos of the *yap1^wt/wt^wwtr1^wt/ncv114^* (left) and the *yap1^ncv117^wwtr1^ncv114^* (right) at 26 hpf. Dotted lines indicate the heart region. Note the CPCs located bilaterally in the *yap1^ncv117^wwtr1^ncv114^* embryos. The confocal 3D-stack images are a set of representative images of three independent experiments.

**Figure 3—Figure supplement 1.**
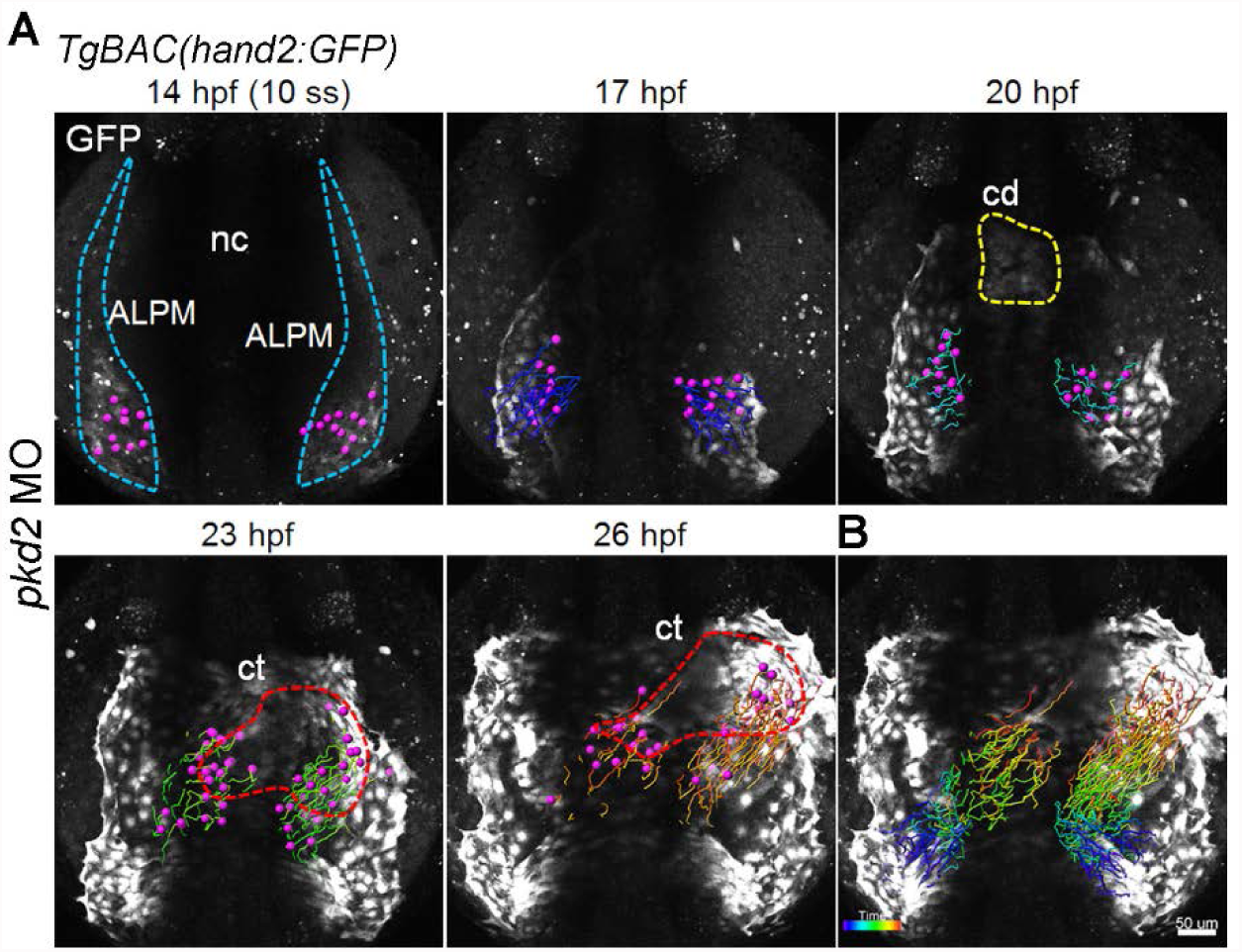
Caudal region of right ALPM migrate toward the venous pole of reversed-heart tube in the *pkd2* morphant. **(A)** Time-sequential 3D-rendered confocal images of a *TgBAC*(*hand2:GFP*) embryo injected with *pkd2* MO from 14 hpf (10 ss) to 26 hpf as indicate at the top. Spots of magenta denote the cells in the caudal part of left and right ALPM. **(B)** Tracking of caudal end *hand2* promoter-activated ALPM cells from 14 hpf to 26 hpf. The color of the tracks changes from blue to red according to the time after imaging (0 h to 12 h). Notochord, nc; cardiac disc, cd; cardiac tube, ct. ALPM, cd, and ct are marked by the blue, yellow, and red broken lines, respectively. The confocal 3D-stack images are a set of representative images of four independent experiments.

**Figure 4—Figure supplement 1.**
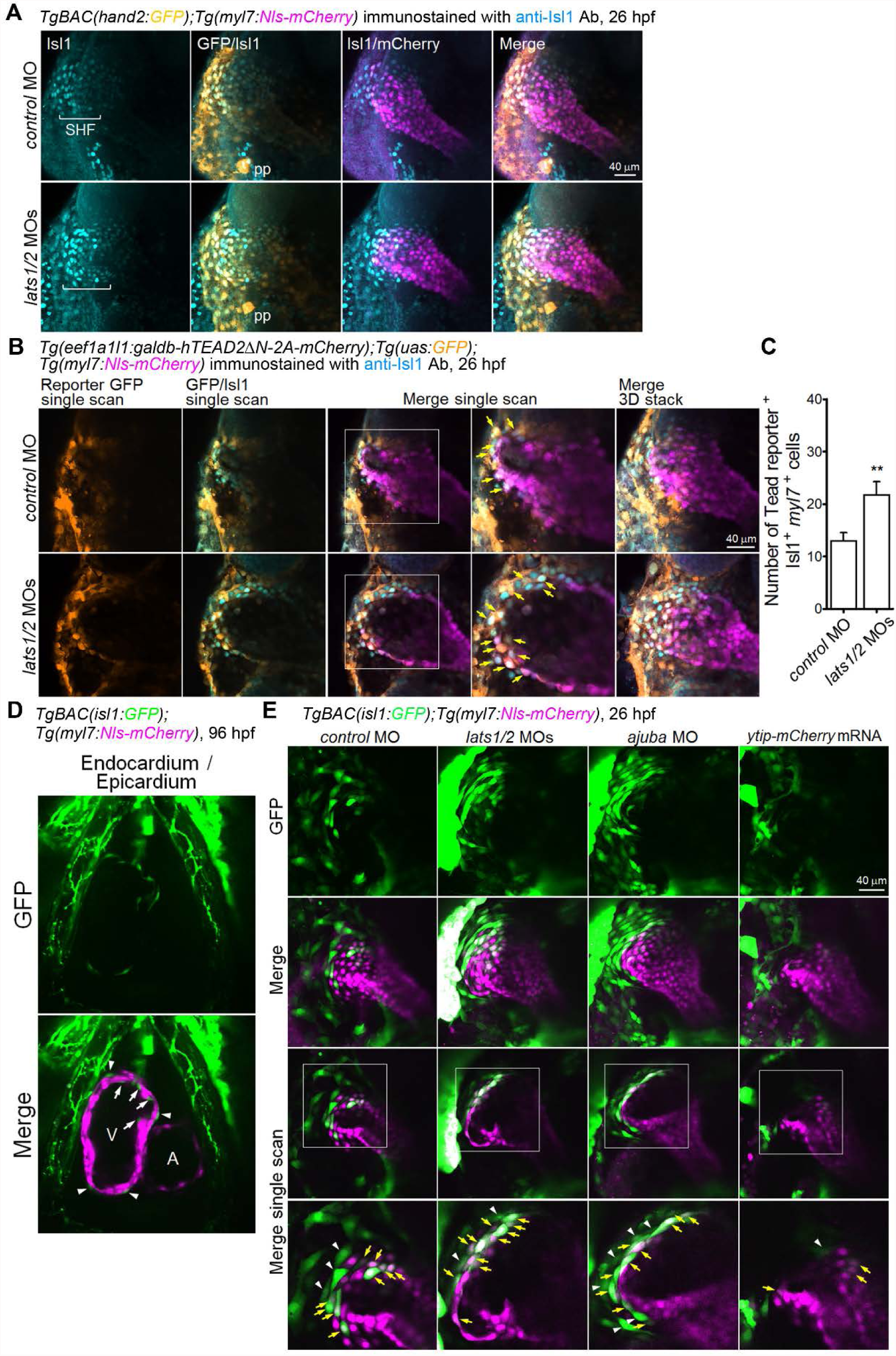
Depletion of Lats1/2 leads to an increase of the number of Isl1-positive SHF cells in the venous pole. **(A)** 3D confocal stack images of *TgBAC*(*hand2:GFP);Tg(myl7:Nls-mCherry*) embryos injected with MO indicated at the left and immunostained with anti-Isl1 antibody (Ab) at 26 hpf. Brackets denote the SHF cells that are Isl1-positive and both *hand2* and *myl7* promoter-activated cells and those in contact with *myl7* promoter-activated cell. The first, second, third and fourth images are Isl-1 immunostaining, the merged image of GFP image and Isl1 immunostaining, the merged image of Isl1 immunostaining and mCherry image, and the merged of all the three (GFP, mCherry, and Isl1 immunostaining), respectively. pp indicates the pharyngeal pouch that expresses *hand2* promoter-activated GFP signals. Dorsal view, anterior to the top. **(B)** Confocal images of *Tg(eef1a1l1:galdb-hTEAD2*Δ*N-2A-mCherry);Tg(uas:GFP*)*;Tg(myl7:Nls-mCherry)* embryos injected with the MO indicated at the left and immunostained with anti-Isl1 Ab at 26 hpf. The boxed regions in the center panels are enlarged in the next right panels. Tead reporter-2 dependent GFP-positive cells in the Isl1-positive *myl7* promoter-activated cells (yellow arrows) are observed in the venous pole. 3D confocal stack images (the most right panels) and single scan images (left four panels). Dorsal view, anterior to the top. **(C)** Quantitative analyses of the number of Isl1-positive *myl7* promoter-activated cells that are positive for Tead reporter-6 dependent GFP in the venous pole of **(B)** (n=5). ^**^p < 0.01. **(D)** Single-scan confocal images of the *TgBAC*(*isl1:GFP);Tg(myl7:Nls-mCherry*) embryos at 96 hpf. GFP-positive mCherry-negative cells in the inside (arrows) and outside (arrowheads) of *myl7* promoter-positive ventricular CMs are endocardial cells and epicardial cells, respectively. A, atrium; V, ventricle. Ventral view, anterior to the top. **(E)** Confocal images of the *TgBAC*(*isl1:GFP);Tg(myl7:Nls-mCherry*) embryos injected with the MO and mRNA at 26 hpf. The boxed regions are enlarged in the bottom panels. Yellow arrows indicate both *isl1* and *myl7* promoter-activated cells in the venous pole. White arrowheads indicate the *isl1* promoter-activated cells in contact with *myl7* promoter-4 activated cells. 3D confocal stack images (upper two panels) and single-scan images (bottom two panels). Dorsal view, anterior to the top. The confocal 3D-stack images and single-scan images are a set of representative images of at least three independent experiments.

**Figure 5—Figure supplement 1.**
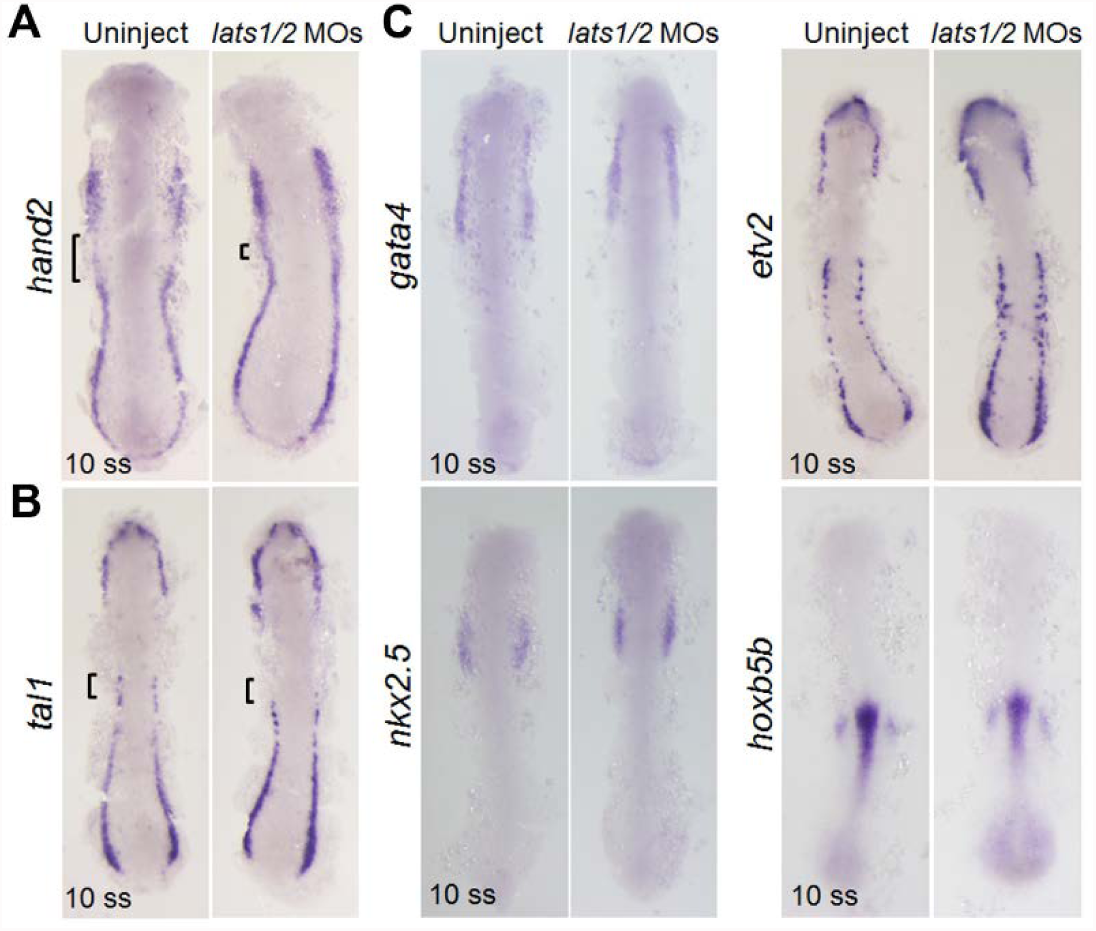
Depletion of Lats1/2 leads to an increase in the expression of *hand2*. **(A-C)** Whole mount in situ hybridization (WISH) analyses of the embryos of the control (uninject, left panels) and injected with *lats1/2* MOs (right panels) using antisense probe indicated at the left side of panels at the 10 somite stage (ss). **(A)** Brackets indicate the gap between *hand2*-positive regions of ALPM and PLPM. Note that the gap is absent in the *lats1/2* morphants. **(B)** Brackets indicate the *tal1*-positive rostral end of PLPM in the control. Note that the region indicated by the bracket in the uninjected embryo is absent in the *lats1/2* morphants. Dorsal view, anterior to the top. WISH images are a set of representative images of four independent experiments.

**Figure 6—Figure supplement 1.**
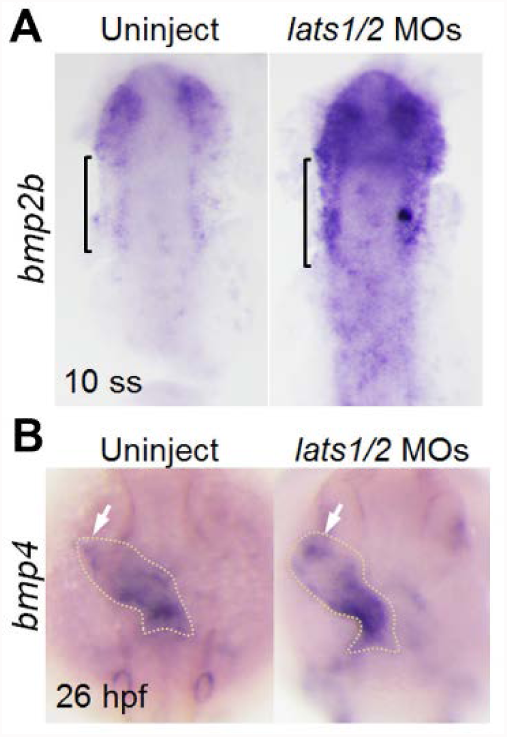
Depletion of Lats1/2 leads to an increase in the *bmps* expression. **(A, B)** WISH analyses of the embryos at 10 ss **(A)** and 26 hpf **(B)** of the control (uninject, left panels) and injected with *lats1/2* MOs (right panels) using antisense probe for *bmp2b* **(A)** and *bmp4* **(B)**. **(A)** Brackets indicate the *bmp2b*-positive left ALPM. **(B)** Arrows indicate the *bmp4*-positive cells in the venous pole. Broken lines indicate the heart tube. WISH images are a set of representative images of three independent experiments.

**Figure 6—Figure supplement 2.**
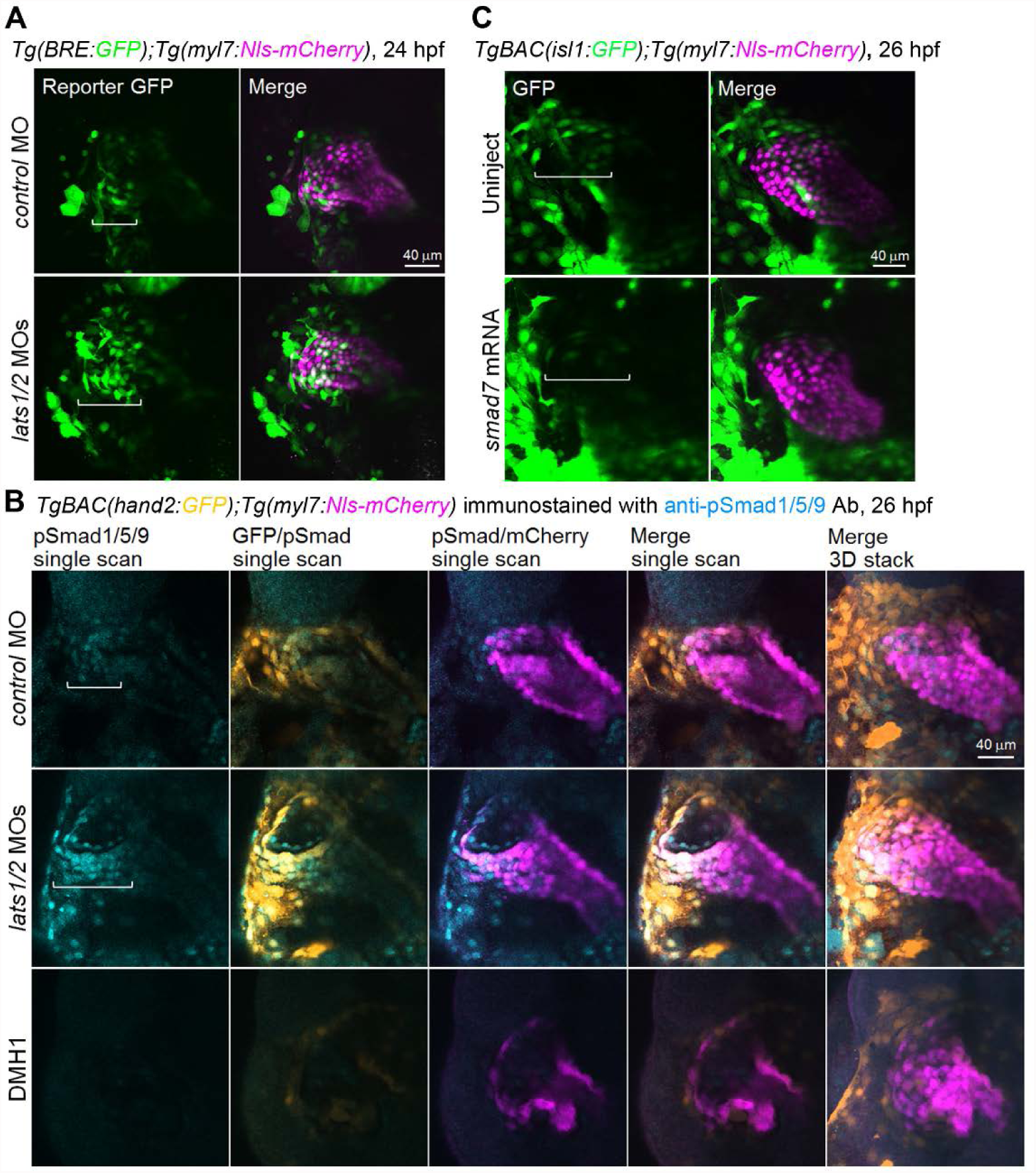
Depletion of Lats1/2 leads to an activation of Bmp-Smad signaling that is necessary for Isl1-positive SHF formation. **(A)** 3D confocal stack images of *Tg*(*BRE:GFP);Tg(myl7:Nls-mCherry*) embryos injected with the *control* MO and *lats1/2* MOs at 24 hpf. Brackets indicate the region of BRE-dependent GFP-positive *myl7* promoter-activated cells in the venous pole. **(B)** Confocal images of the *TgBAC(hand2:GFP);Tg(myl7:Nls-mCherry)* embryos injected with the *control* MO, *lats1/2* MOs, followed by the treatment with DMH1 (10 μM) from 14 hpf to 26 hpf (upper to bottom panels) and immunostained with anti-pSmad1/5/9 Ab at 26 hpf. Brackets indicate the phosphorylated Smad1/5/9-positive *myl7* promoter-activated cells in the venous pole. 3D confocal stack images (the most right panels) and single scan images (left four panels). Note that both *hand2* promoter-activated and pSmad1/5/9-positive cells in the venous pole are decreased in the embryos treated with DMH1. Dorsal view, anterior to the top. **(C)** 3D confocal stack images of *TgBAC*(*isl1:GFP);Tg(myl7:Nls-mCherry*) embryos of the control (uninject, upper panels) and injected with 100 pg *smad7* mRNA (bottom panels) at 26 hpf. Brackets indicate the region of *isl1* promoter-activated GFP-positive SHF cells in the venous pole. The confocal 3D-stack images and single-scan images are a set of representative images of at least three independent experiments.

## Legends for videos

**Video 1.** *hand2* promoter-activated cells in the caudal region of the left ALPM move to venous pole. Time lapse recording of 3D-rendered confocal images of a *TgBAC*(*hand2:GFP*) embryo from 14 hpf (10 ss) to 26 hpf. Note the migration of caudal region of left (magenta) and right (cyan) ALPM cells toward venous pole and arterial pole of heart tube, respectively. Changes of colors reflect the tracking time (blue, 0 h; red, 12 h). Dorsal view, anterior to the top. The time lapse movie is a set of representative data of six independent experiments. Video 1 is related to Figure 3A, B.

**Video 2.** In the reversed-heart, *hand2* promoter-activated cells in the caudal region of the right ALPM move to venous pole. Time lapse recording of 3D-rendered confocal images of a *TgBAC*(*hand2:GFP*) embryo injected with *pkd2* MO from 14 hpf (10 ss) to 26 hpf. Note the migration of caudal region of left and right ALPM cells (magenta) was reversed. Changes of colors reflect the tracking time (blue, 0 h; red, 12 h). Dorsal view, anterior to the top. The time lapse movie is a set of representative data of four independent experiments. Video 2 is related to Figure 3A, B.

**Video 3.** Tead reporter-activated cells in the caudal region of the left ALPM move to venous pole. Time lapse recording of 3D-rendered confocal images of a *Tg(eef1a1l1:galdb-hTEAD2*Δ*N-2A-mCherry);Tg(uas:GFP*) embryo from 14 hpf (10 ss) to 24 hpf. Note the migration of Tead reporter-activated cells (magenta) in the caudal region of the left ALPM toward the venous pole of heart tube. Changes of colors reflect the tracking time (blue, 0 h; red, 10 h). Dorsal view, anterior to the top. The time lapse movie is a set of representative data of six independent experiments. Video 3 is related to Figure 3D, E.

**Video 4.** Bmp-Smad signal-activated cells in the caudal region of the left ALPM move to venous pole. Time lapse recording of 3D-rendered confocal images of a *Tg(BRE:GFP*)*;Tg(myl7:Nls-mCherry)* embryo from 14 hpf (10 ss) to 24 hpf. Note that Bmp-Smad signal is activated in the caudal region of the left ALPM (cyan) and that those cells move to venous pole of heart tube. Changes of colors reflect the tracking time (blue, 0 h; red, 10 h). Dorsal view, anterior to the top. The time lapse movie is a set of representative data of six independent experiments. Video 4 is related to Figure 6A.

